# The Stochastic Early Reaction, Inhibition, and Late Action (SERIA) Model for Antisaccades

**DOI:** 10.1101/109090

**Authors:** Eduardo A. Aponte, Dario Schoebi, Klaas E. Stephan, Jakob Heinzle

## Abstract

The antisaccade task is a classic paradigm used to study the voluntary control of eye movements. It requires participants to suppress a reactive eye movement to a visual target and to concurrently initiate a saccade in the opposite direction. Although several models have been proposed to explain error rates and reaction times in this task, no formal model comparison has yet been performed. Here, we describe a Bayesian modeling approach to the antisaccade task that allows us to formally compare different models on the basis of their evidence. First, we provide a formal likelihood function of actions (pro- and antisaccades) and reaction times based on previously published models. Second, we introduce the *Stochastic Early Reaction, Inhibition, and late Action model* (SERIA), a novel model postulating two different mechanisms that interact in the antisaccade task: an early GO/NO-GO race decision process and a late GO/GO decision process. Third, we apply these models to a data set from an experiment with three mixed blocks of pro- and antisaccade trials. Bayesian model comparison demonstrates that the SERIA model explains the data better than competing models that do not incorporate a late decision process. Moreover, we show that the race decision processes postulated by the SERIA model are, to a large extent, insensitive to the cue presented on a single trial. Finally, we use parameter estimates to demonstrate that changes in reaction time and error rate due to the probability of a trial type (prosaccade or antisaccade) are best explained by faster or slower inhibition and the probability of generating late voluntary prosaccades.

**Author summary:** One widely replicated finding in schizophrenia research is that patients tend to make more errors in the antisaccade task, a psychometric paradigm in which participants are required to look in the opposite direction of a visual cue. This deficit has been suggested to be an endophenotype of schizophrenia, as first order relatives of patients tend to show similar but milder deficits. Currently, most models applied to experimental findings in this task are limited to fit average reaction times and error rates. Here, we propose a novel statistical model that fits experimental data from the antisaccade task, beyond summary statistics. The model is inspired by the hypothesis that antisaccades are the result of several competing decision processes that interact nonlinearly with each other. In applying this model to a relatively large experimental data set, we show that mean reaction times and error rates do not fully reflect the complexity of the processes that are likely to underlie experimental findings. In the future, our model could help to understand the nature of the deficits observed in schizophrenia by providing a statistical tool to study their biological underpinnings.

## Introduction

In the antisaccade task ([1]; for reviews, see [2,3]), participants are required to saccade in the contralateral direction of a visual cue. This behavior is thought to require both the inhibition of a reflexive saccadic response towards the cue and the initiation of a voluntary eye movement in the opposite direction. A failure to inhibit the reflexive response leads to an erroneous saccade towards the cue (i.e., a prosaccade), which is often followed by a corrective eye movement in the opposite direction (i.e., an antisaccade). As a probe of inhibitory capacity, the antisaccade task has been widely used to study psychiatric and neurological diseases [3]. Notably, since the initial report [4], studies have consistently found an increased number of errors in patients with schizophrenia when compared to healthy controls, independent of medication and clinical status [5-8]. Moreover, there is evidence that an increased error rate constitutes an endophenotype of schizophrenia, as antisaccade deficits are also present in non-affected, first-degree relatives of diagnosed individuals (for example [5,7]; but for negative findings see for example [9][10]).

Unfortunately, the exact nature of the antisaccade deficits and their biological origin in schizophrenia remain unclear. One path to improve our understanding of these experimental findings is to develop generative models of their putative computational and/or neurophysiological causes [11]. Generative models that capture the entire distribution of responses can reveal features of the data that are not apparent when only considering summary statistics such as mean error rate (ER) and reaction time (RT) [12-15]. Additionally, this type of model can potentially relate behavioral findings in humans to their biological substrate.

Here, we apply a generative modeling approach to the antisaccade task. First, we introduce a novel model of this paradigm based on previous proposals [16-20]. For this, we formalize the ideas introduced by Noorani and Carpenter [17] and extend them into what we refer to as the *Stochastic Early Response, Inhibition and late Action* (SERIA) model. Second, we apply both models to an experimental data set of three mixed blocks of pro- and antisaccades trials with different trial type probability using formal Bayesian inference. More specifically, we compare several models using Bayesian model comparison. Third, we use the parameter estimates from the best model to investigate the effects of our experimental manipulation. We found that there was positive evidence in favor of the SERIA model when compared to our formalization of the model proposed in [17]. Moreover, the parameters estimated through model inversion revealed the complexity of the decision processes underlying the antisaccade task that is not obvious from mean RT and ER.

This paper is organized as follows. First, we formalize the model developed in [17] and introduce the SERIA model. Second, we describe our experimental setup. Third, we present our behavioral findings in terms of summary statistics (mean RT and ER), the comparison between different models, and the parameter estimates. Finally, we review our findings, discuss other recent models, potential future developments, and translational applications.

## Materials and methods

### Ethics statement

All participants gave written informed consent before the study. All experimental procedures were approved by the local ethics board (Kantonale Ethikkomission Zürich, KEK-ZH-Nr.2014-0246).

### Race models for antisaccades

In this section, we derive a formal description of the models evaluated in this paper. We start with a formalized version of the model proposed by Noorani and Carpenter in [17] and proceed to extend it. Their approach resembles the model originally proposed by Camalier and colleagues [21] to explain RT and ER in the double step and search step tasks, in which participants are either asked to saccade to successively presented targets or to saccade to a target after a distractor was shown. Common to all these tasks is that subjects are required to inhibit a prepotent reaction to an initial stimulus and then to generate an action towards a secondary goal. Briefly, Camalier and colleagues [21] extended the original ‘horse-race’ model [16] by including a secondary action in countermanding tasks. In [17], Noorani and Carpenter used a similar model in combination with the LATER model [22] in the context of the antisaccade task by postulating an endogenously generated inhibitory signal. Note that this model, or variants of it, have been used in several experimental paradigms (reviewed in [20]). Here, we limit our discussion to the antisaccade task.

#### The pro, stop, and antisaccade model (PROSA)

Following [17], we assume that the RT and the type of saccade generated in a given trial are caused by the interaction of three competing processes or units. The first unit *u_p_* represents a command to perform a prosaccade, the second unit *u_s_* represents an inhibitory command to stop a prosaccade, and the third unit *u_a_* represents a command to perform an antisaccade. The time *t* required for unit *u_i_* to arrive at threshold *s_i_* is given by:

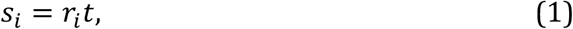

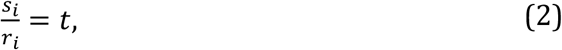

where *r_i_* represents the slope or increase rate of unit *u_i_*, *s_i_* represents the height of the threshold, and *t* represents time. We assume that, on each trial, the increase rates are stochastic and independent of each other.

The time and order in which the units reach their thresholds *s_i_* determines the action and RT in a trial. If the prosaccade unit *u_p_* reaches threshold before any other unit at time *t*, a prosaccade is elicited at *t*. If the antisaccade unit arrives first, an antisaccade is elicited at *t*. Finally, if the stop unit arrives before the prosaccade unit, an antisaccade is elicited at the time when the antisaccade unit reaches threshold. It is worth mentioning that, although this model is motivated as a race-to-threshold model, actions and RTs depend only on the arrival times of each of the units and ultimately no explicit model of increase rates or thresholds is required. Thus, for the sake of clarity, we refer to this approach as a ‘race’ model, in contrast to ‘race-to-threshold’ models that explicitly describe increase rates and thresholds.

Formally (but in a slight abuse of language), the two random variables of interest, the reaction time *T* ∈ [0, ∞[ and the type of action performed *A* ∈ {*pro*, *anti*}, depend only on three further random variables: the arrival times *U_p_*, *U_s_*, *U_a_* ∈ [0, ∞[ of each of the units. The probability of performing a prosaccade at time *t* is given by the probability of the prosaccade unit arriving at time *t*, and the stop and antisaccade unit arriving afterwards:

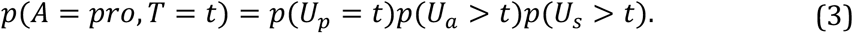

The probability of performing an antisaccade at time *t* is given by

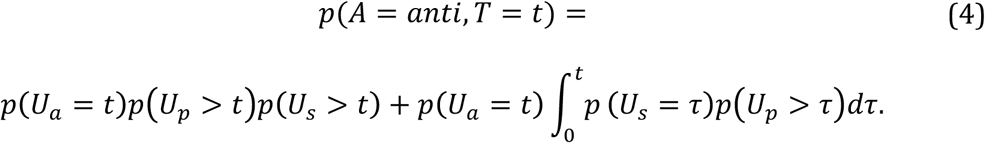

The first term on the right side of Eq. 4 corresponds to the unlikely case that the antisaccade unit arrives before the prosaccade and the stop unit. The second term describes trials in which the stop unit arrives before the prosaccade unit. It can be decomposed into two terms:

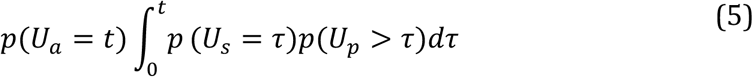

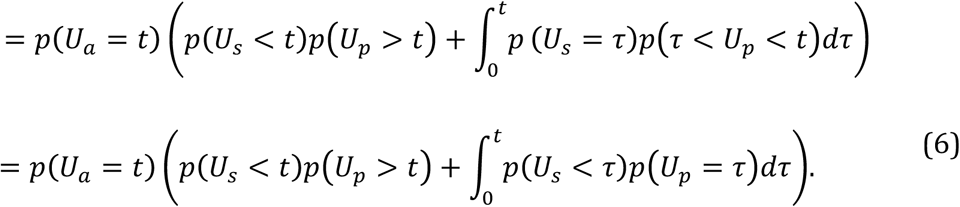

The term 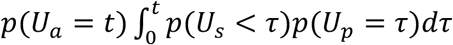 describes the condition in which the prosaccade unit is inhibited by the stop unit allowing for an antisaccade. Note that if the prosaccade unit arrives later than the antisaccade unit, the arrival time of the stop unit is irrelevant. That means that we can simplify Eq. 4 to

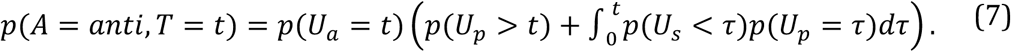

Eq. 3 and 7 constitute the likelihood function of a single trial, defining the joint probability of an action and the corresponding RT. We refer to this likelihood function as the PROStop-Antisaccade (PROSA) model. It shares the central assumptions of [17], namely: (i) the time to reach threshold of each of the units is assumed to depend linearly on the rate *r*, (ii) it includes a stop unit whose function is to inhibit prosaccades and (iii) there is no lateral inhibition between the different units. Finally, (iv) RTs are assumed to be equal to the arrive-at-threshold times. Note that the RT distributions are different from the arrival time distributions because of the interactions between the units described above. The main difference of this model compared to [17] is that we do not exclude *a priori* the possibility of the antisaccade unit arriving earlier than the other units. Aside from this, both models are conceptually equivalent.

### The Stochastic Early Reaction, Inhibition, and Late Action Model (SERIA)

The PROSA model is characterized by a strict association between units and action types. In other words, the unit *u_p_* leads unequivocally to a prosaccade, whereas the unit *u_a_* always triggers an antisaccade. This implies that if the distribution of the arrival times of the units is unimodal and strictly positive, the PROSA model cannot predict voluntary slow prosaccades with a late peak, or in simple words, the PROSA model cannot account for slow, voluntary prosaccades that have been postulated in the antisaccade task [23]. Similarly, it has been argued that prosaccade RT can be described by the mixture of two distributions (for example [2,22]). To account for this, we introduce the Stochastic Early Reaction, Inhibition and Late Action (SERIA) model.

According to this model, and in analogy to the PROSA model, an early reaction takes place at time *t* if the early unit *u_e_* arrives before the late and inhibitory units, *u_l_* and *u_i_*, respectively. If the inhibitory or late unit arrives before the early unit, a late response is triggered at the time the late unit reaches threshold. Crucially, both early and late responses can trigger pro- and antisaccades with a certain probability. Thus, in parallel to the race processes which determine RTs, an independent, secondary decision process is responsible for which reaction is generated. Fig. 1 shows the structure of the SERIA model.

**Fig 1.**
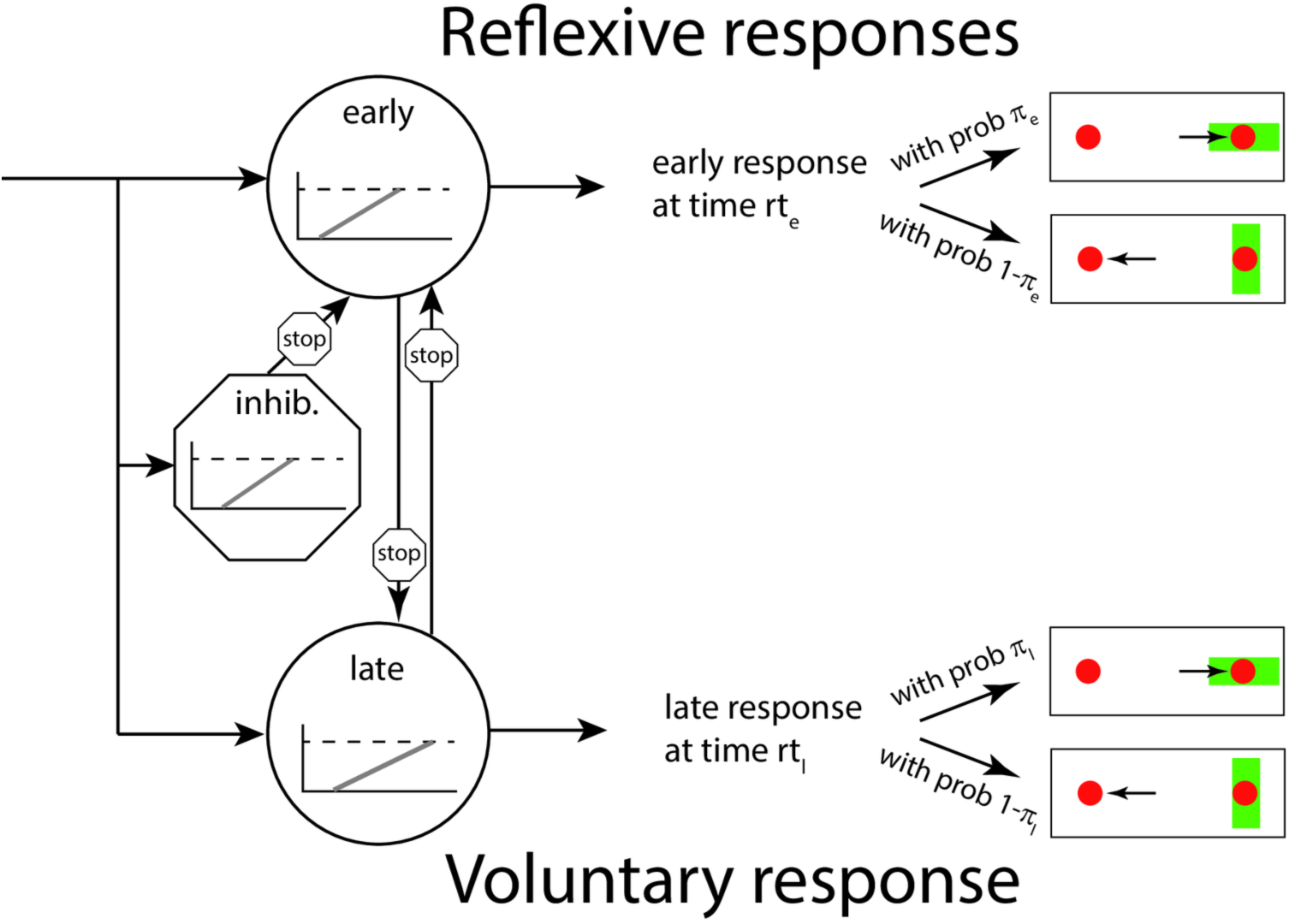
Layout of the SERIA model. The presentation of a visual cue (a green bar) triggers the race of three independent units. The inhibitory unit can stop an early response. Importantly, both early and late responses can trigger pro- and antisaccades. Note that the PROSA model is a special case of the SERIA model in which *π_e_* = 1 and *π_l_* = 0, i.e. all early responses are prosaccades, whereas all late responses are antisaccades.

To formalize the concept of early and late responses, we introduce a new unobservable random variable that represents the type of response *R* ∈ {*early, late*}. The distribution of the RTs is analogous to the PROSA-model, such that, for instance, the probability of an early response at time *t* is given by

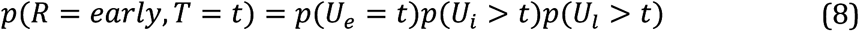

where *U_e_, U_i_*, and *U_l_* represent the arrival times of the early, inhibitory, and late units respectively. The fundamental assumption of the SERIA model is that a secondary decision process, beyond the race between early, inhibitory, and late units, decides the action generated in a single trial. An initial approach to model this secondary decision process is to assume that the action type (pro- or antisaccade) is conditionally independent of the RT given the response type (early or late). Hence, the distribution of RTs is not *a priori* coupled to the saccade type anymore; RT distributions for both pro- and antisaccades could in principle be bimodal, consisting of both fast reactive and slow voluntary saccades.

Formally, the conditional independency assumption can be written down as

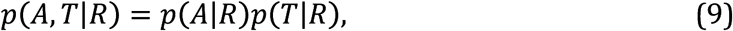

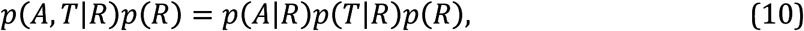

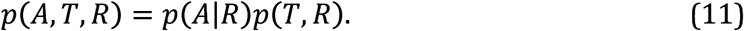

The term *p*(*A*|*R*) is simply the probability of an action, given a response type. We denote it as

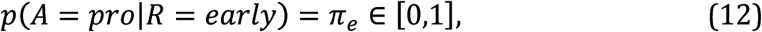

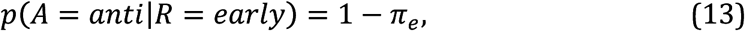

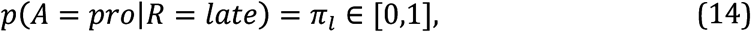

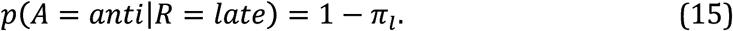

Since the type of response *R* is not observable, it is necessary to marginalize it out in Eq. [11] to obtain the likelihood of the SERIA model:

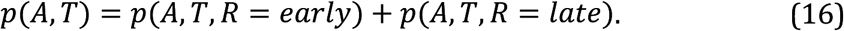

The complete likelihood of the model is given by substituting the terms in Eq. [16]:

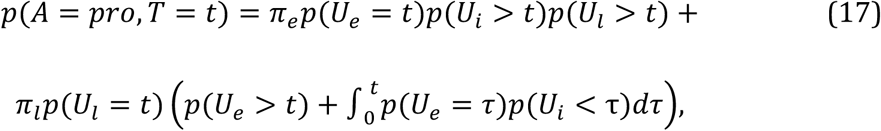

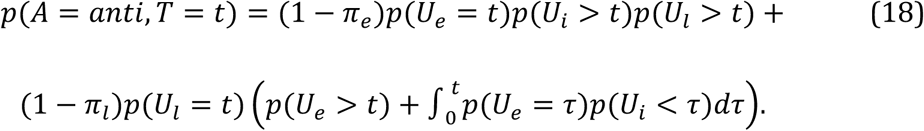

It is worth noting here that the PROSA model is a special case of the SERIA model, namely, it corresponds to the assumption that *π_e_* = 1 and *π_l_* = 0. The SERIA model allows for bimodal distributions, as both early and late responses can be pro- and antisaccades. Importantly, one prediction of the model is that late prosaccades have the same distribution as late antisaccades.

### Late race competition model for saccade type

Until now, we have assumed that the competition that leads to late pro- and antisaccades does not depend on time in the sense that late actions are conditionally independent of RT. This assumption can be weakened by postulating a secondary race between late responses; this leads us to a modified version of the SERIA model, that we refer to as the late race SERIA model (SERIA_lr_). The derivation proceeds similarly to the SERIA model, except that we postulate a fourth unit that generates late prosaccades instead of assuming that the late decision process is time insensitive.

This version of the SERIA model includes an early unit *u_e_* that, for simplicity, we assume produces only prosaccades, an inhibitory unit that stops early responses *u_i_*, a late unit that triggers antisaccades *u_a_*, and a further unit that triggers late prosaccades *u_p_*. As before, if the early unit reaches threshold before any other unit, a prosaccade is generated with probability

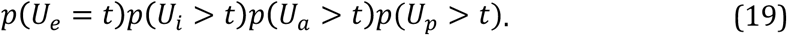

If any of the late units arrive first, the respective action is generated with probability:

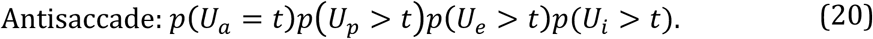

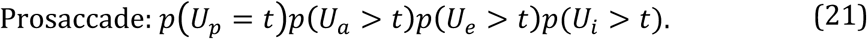

Finally, if the inhibitory unit arrives first, either a late pro- or antisaccade is generated with probability

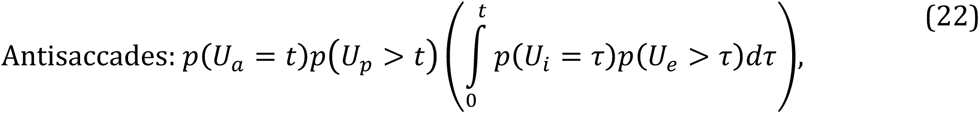

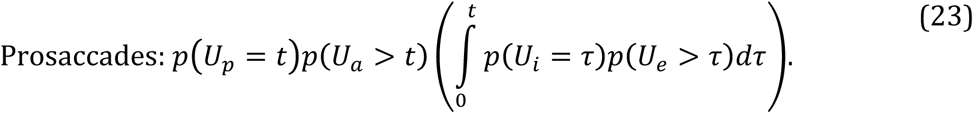

Implicit in the last two terms is the competition between the late units, which are assumed again to be independent of each other. Formally, this competition is expressed as the probability of, for example, the late antisaccade unit arriving before a late prosaccade *p*(*U_a_* = *t*)*p*(*U_p_* > *t*). A schematic representation of the model is shown in Fig. 2. This late race is similar to the Linear Ballistic Accumulation model proposed by [24], although in that model decisions are seen as the result of a race of ballistic accumulation processes with fixed threshold but stochastic base line and increase rate. Here we only assume that the late decision process is a GO-GO race [21].

**Fig 2.**
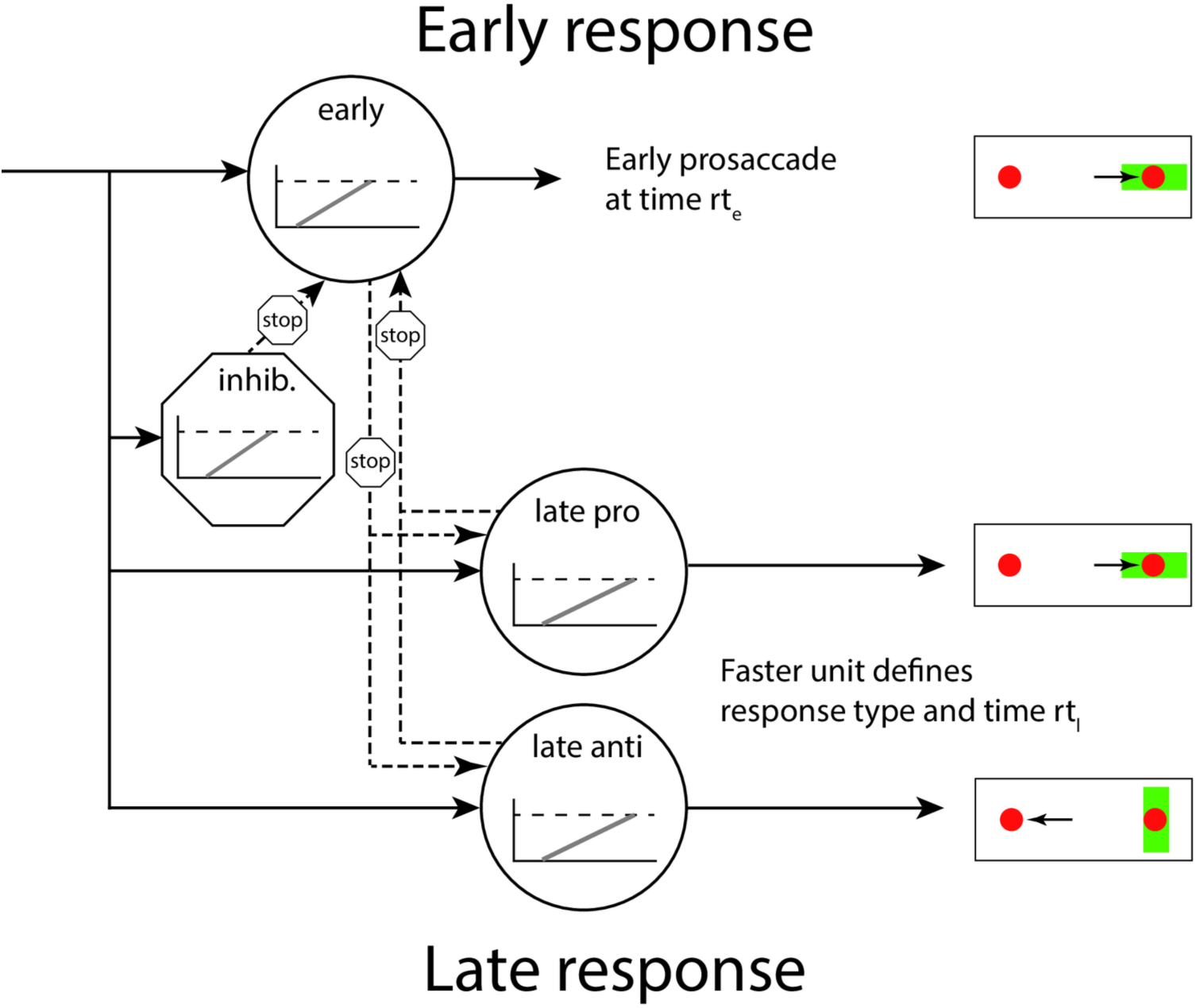
Layout of the SERIA_lr_ model. The presentation of a visual cue (a green bar) triggers the race of four independent units. The inhibitory unit can stop an early response. The late decision process is triggered by the competition between two further units.

The likelihood of an action is given by summing over all possible outcomes that lead to that action:

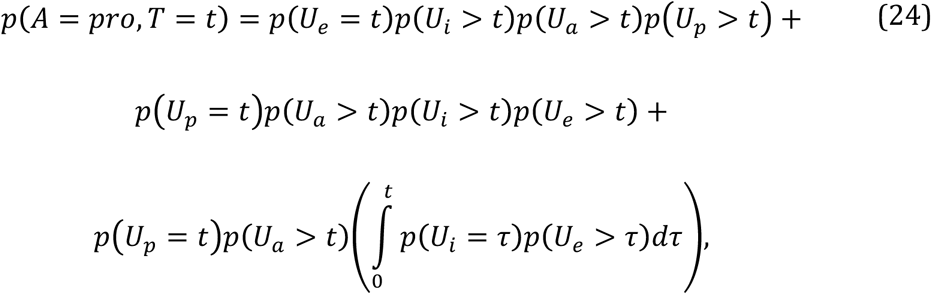

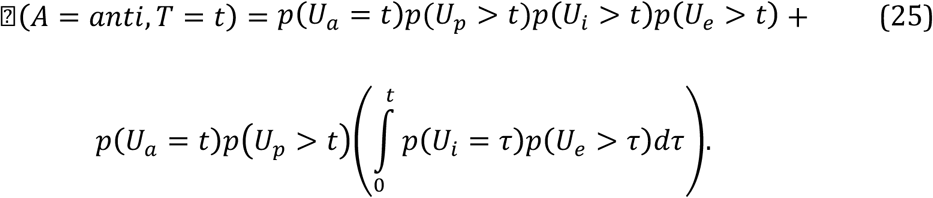

We have left out some possible simplifications in Eq. 24 and 25 for the sake of clarity.

The conditional probability of a late antisaccade is given by the interaction between the late units, such that

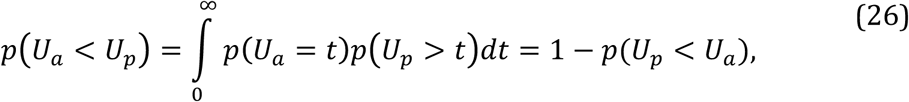

is analogous to the probability of a late antisaccade 1 − *π_l_* in the SERIA model. This observation shows that the main difference between the SERIA and SERIA_lr_ model is that the former postulates that the distribution of late pro- and antisaccades are equal and conditionally independent of the action performed, whereas the latter constrains the probability of a late antisaccade to be a function of the arrival times of the late units.

The expected *response time* of late pro and antisaccade actions is given by

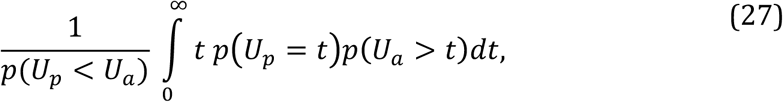

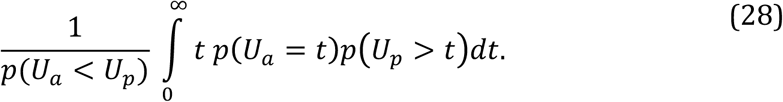

We will refer to these terms as the mean *response time* of pro- and antisaccade actions, in contrast to the mean arrival times, which are the expected value of any single unit.

### Non-decision time

The models above can be further finessed to account for non-decision times *δ* by transforming the RT *t* to *t_δ_* = *t* − *δ*. The delay *δ* might be caused by, for example, conductance delays from the retina to the cortex. In addition, the antisaccade or late units might include a constant delay *δ_a_*, which is often referred to as the antisaccade cost [1]. Note that the model is highly sensitive to *δ* because any RT below it has zero probability. In order to relax this condition and to account for early outliers, we assumed that saccades could be generated before *δ* at a rate *η* ∈ [0,1] such that the marginal likelihood of an outlier is

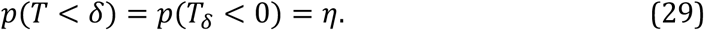

For simplicity, we assume that outliers are generated with uniform probability in the interval [0,*δ*]:

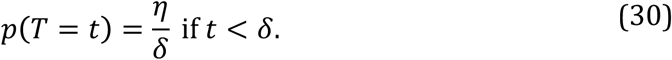

Furthermore, we assume that the probability of an early outlier being a prosaccade was approximately 100 times higher than being an antisaccade. Because of the new parameter *η*, the distribution of saccades with a RT larger than *δ* needs to be renormalized by the factor 1 − *η*. In the case of the PROSA model, for example, this means that the joint distribution of action and RT is given by the conditional probability

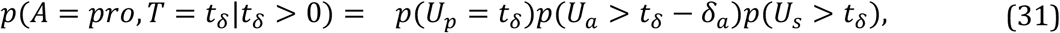

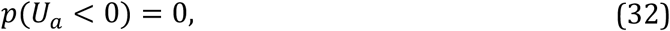

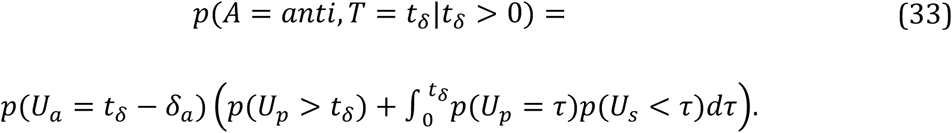

A similar expression holds for the SERIA models. However, in the PROSA model a unit-specific delay is equal to an action-specific delay. By contrast, in the SERIA model both early and late responses can generate pro- and antisaccades. Thus, *δ_a_*, represents a delay in the late actions that affects both late pro- and antisaccades.

### Parametric distributions of the increase rate

The models discussed in the previous sections can be defined independently of the distribution of the rate of each of the units. In order to fit experimental data, we considered four parametric distributions with positive support for the rates: gamma [13], inverse gamma, lognormal [25] and the truncated normal distribution (similarly to [22] and [24]). Table 1 and Fig 3 summarize these distributions, their parameters, and the corresponding arrival time densities. We considered five different configurations: 1) all units were assigned *inverse gamma* distributed rates, 2) all units were assigned *gamma* distributed rates, 3) the increase rate of the pro and stop units (or early and the inhibitory units) was *gamma distributed* but the antisaccade (late) unit’s increase rate was *inverse gamma* distributed, 4) all the units were assigned *lognormal* distributed rates or 5) all units were assigned *truncated normal* distributed rates.

**Table 1:** Parametric density functions of the increase rates.

| Name | Parameters | Rate p.d.f. | Arrival time p.d.f. |
| --- | --- | --- | --- |
| Gamma | $k, \theta$ | $\frac{\theta^{-k}}{\Gamma(k)} e^{-r/\theta} r^{k-1}$ | $\frac{\theta^k}{\Gamma(k)} e^{-\theta/t} t^{-k-1}$ |
| Inv. gamma | $k, \theta$ | $\frac{\theta^k}{\Gamma(k)} e^{-\theta/r} r^{-k-1}$ | $\frac{\theta^{-k}}{\Gamma(k)} e^{-t/\theta} t^{k-1}$ |
| Log normal | $\mu, \sigma^2$ | $\frac{1}{\sqrt{2\pi}\sigma r} e^{-\frac{1}{2}\left(\frac{\ln r - \mu}{\sigma}\right)^2}$ | $\frac{1}{\sqrt{2\pi}\sigma t} e^{-\frac{1}{2}\left(\frac{\ln t + \mu}{\sigma}\right)^2}$ |
| T. normal | $\mu, \sigma^2$ | $\frac{1}{Z} e^{-\frac{1}{2}\left(\frac{r - \mu}{\sigma}\right)^2}$ | $\frac{1}{Z t^2} e^{-\frac{1}{2}\left(\frac{t^{-1} - \mu}{\sigma}\right)^2}$ |

Z is the normalization constant $Z = \int_0^\infty \exp\left(-\frac{(r-\mu)^2}{2\sigma^2}\right) dr$ .

**Fig 3.**
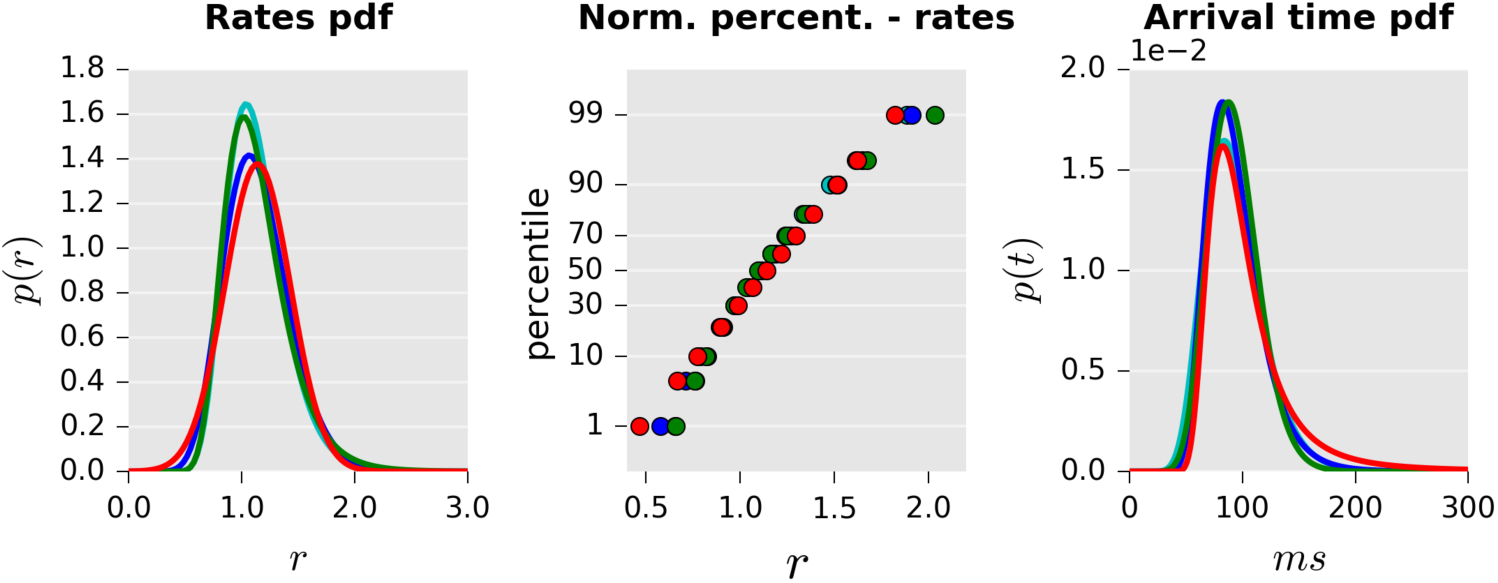
Illustration of probability distributions used to model increase rates. Left: Distribution of the rates based on different probability density functions: Normal (red), gamma (blue), inverse gamma (green), and log-normal (cyan). All distributions were matched to have equal mean and variance. Center: Probit plots of the same distributions. While the gamma and lognormal distributions are very close to the straight line induced by the normal distribution, the inverse gamma distribution diverges slightly more from linearity. Right: Arrival time distribution (scaled to ms).

All the parametric distributions considered here can be fully characterized by two parameters which we generically refer to as *k* and *θ*. Hence, the PROSA model is characterized by the parameters for each unit *k_p_, k_a_, k_s_,θ_p_, θ_a_, θ_S_*. The SERIA model can be characterized by analogous parameters *k_e_, k_l_, k_i_, θ_e_, θ_l_,θ_i_* and the probabilities of early and late prosaccades *π_e_* and *π_l_*. In the case of the SERIA_lr_ model, the probability of a late prosaccade is replaced by the parameters of a late prosaccade unit *k_p_, θ_p_*. In addition to the unit parameters, all models included the non-decision time *δ* the antisaccade (or late unit) cost *δ_a_*, and the marginal rate of early outliers *η*

### Experimental procedures

In this section, we describe the experimental procedures, statistical methods, and inference scheme used to invert the models above. The data is from the placebo condition of a larger pharmacological study that will be reported elsewhere.

#### Participants

Fifty-two healthy adult males naïve to the antisaccade task were invited to a screening session through the recruitment system of the Laboratory of Social and Neural Systems Research of the University of Zurich. During screening, and after being debriefed about the experiment, subjects underwent an electrocardiogram, a health survey, a visual acuity test, and a color blindness test. Subjects were excluded if any of the following criteria were met: age below 18 or above 40 years, regular smoking, alcohol consumption the day before the experiment, any possible interaction between current medication and levodopa or benserazide, pulse outside the range 55-100bpm, recreational drug intake in the past 6 months, history of serious mental or neurological illness, or if the medical doctor supervising the experiment deemed the participant not apt. All subjects gave their written informed consent to participate in the study and received monetary compensation.

#### Procedure

Each subject was invited to two sessions. During both visits, the same experimental protocol was followed. After arrival, placebo or levodopa (Madopar^®^ DR 250, 200mg of levopa + 50 mg benserazide) was orally administered in the form of shape- and color-matched capsules. The present study is restricted to data from the session in which subjects received placebo. Participants and experimenters were not informed about the identity of the substance. Immediately afterwards subjects were introduced to the experimental setup and to the task through a written document. This was followed by a short training block (see below).

The experiment started 70 minutes after substance administration. Subjects participated in three blocks of 192 randomly interleaved pro- and antisaccade trials. The percentages of prosaccade trials in the three blocks were 20%, 50%, or 80%. This yielded three *prosaccade probability* (PP) conditions: PP20, PP50, and PP80. Thus, in the PP20 block, subjects were presented a prosaccade cue in 38 trials, while in all other 154 trials an antisaccade cue was shown. The order of the trials was randomized in each block, but the same order was used in all subjects and sessions. The order of the conditions was counterbalanced across subjects.

#### Stimulus and apparatus

During the experiment, subjects sat in front of a CRT monitor (Philipps 20B40, distance eye-screen: ≃60*cm*, refresh rate: 75Hz). The screen subtended a horizontal visual angle of 38 degrees of visual angle (dva). Eye movements were recorded using a remote infrared camera (Eyelink II, SR-Research, Canada). Participants’ head was stabilized with a chin rest. Data were stored at a sampling rate of 500 Hz.

During the task, two red dots (0.25dva) that constituted the saccadic targets were constantly displayed at an eccentricity of ±12dva. Displaying the saccadic target before the execution of an antisaccade has been reported to affect saccadic velocity and accuracy, but not RTs [26], and arguably decreases the need for sensorimotor transformations [27]. At the beginning of each trial, a gray fixation cross (0.6x0.6 *dva)* was displayed at the center of the screen. After a random fixation interval (500 to 1000 ms), the cross disappeared, and the cue instructing either a pro- or an antisaccade trial (see below) was shown centered on either of the red dots. As mentioned above, in each block, subjects were presented with a prosaccade cue in either 20, 50, or 80 percent of the trials. The order of the presentation of the cues was randomized. The cue was a green rectangle (3.48x0.8*dva*) displayed for 500ms in either horizontal (prosaccade) or vertical orientation (antisaccade). Once the cue was removed and after 1000ms, the next trial started.

Subjects were instructed to saccade in the direction of the cue when a horizontal bar was presented (prosaccade trial) and to saccade in the opposite direction when a vertical bar was displayed (antisaccade trial, see Fig. 3). See [28,29] for similar task designs.

**Fig 4.**
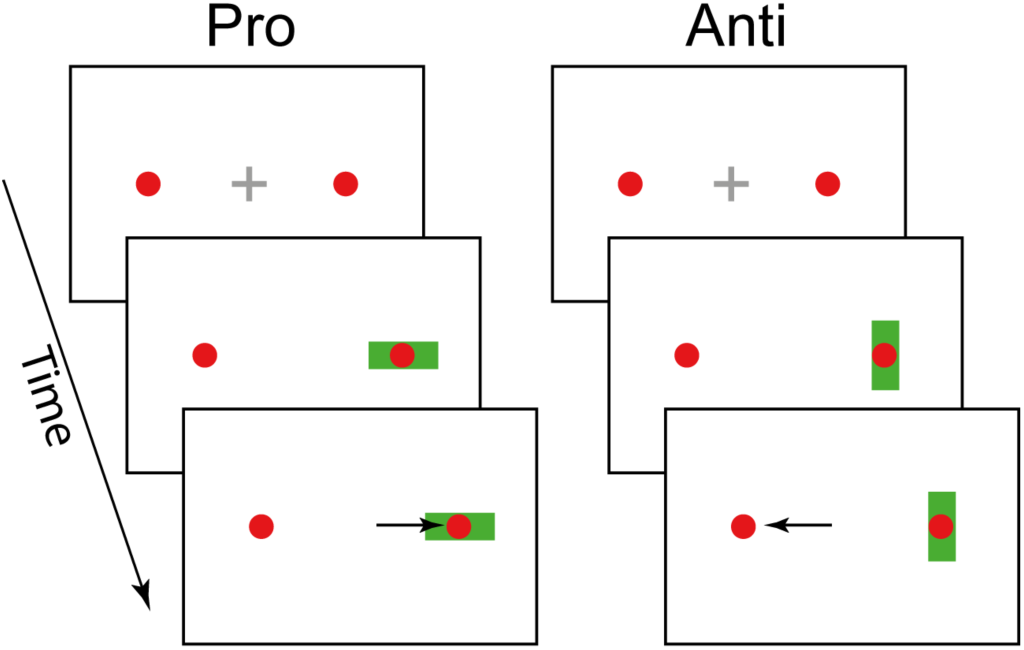
Task design. After a variable fixation period of 500-1000ms (top) the cue (green rectangle) appeared on the screen for 500 ms. The orientation of the cue (horizontal or vertical) indicated the required response (prosaccade or antisaccade).

Prior to the main experiment, participants were trained on the task in a block of 50 prosaccade trials, immediately followed by 50 antisaccade trials. During the training, subjects were automatically informed after each trial whether their response was correct or not (see below), or whether they had failed to produce a saccade within 500ms after cue presentation (CP). No feedback was given during the main experimental blocks.

#### Data preparation

Data were parsed and preprocessed using the Python programming language (2.7). Saccades were detected using the algorithm provided by the eyetracker manufacturer (SR Research), which uses a velocity and acceleration threshold of 22*dva/s* and 3800*dva/s*^2^ [30]. We only considered saccades with a magnitude larger than 2*dva*. RT was defined as the time between CP and the first saccade larger than 2*dva*. A prosaccade trial was considered correct if the end position of the saccade was ipsilateral to the cue and, conversely, an antisaccade trial was considered correct if the end position of the saccade was contralateral to the cue.

Trials were excluded from further analysis if a) data were missing, b) a blink occurred between CP and the main saccade, c) the trial was aborted by the experimenter, d) subjects failed to fixate in the interval between fixation detection and CP, e) if a saccade was detected only later than 800ms after CP, f) if the RT was below 50ms, and in the case of an antisaccade if it was below 110ms. Corrective antisaccades were defined as saccades that a) followed a prosaccade error, b) occurred no later than 900ms after CP, and c) had less than 3*dva* horizontal error from the red circle contralateral to the cue.

Besides the fitted non-decision time *δ* we assumed a fixed non-decision time of 50*ms* for all participants [17]. This was implemented by subtracting 50*ms* of all saccades before being entered into the model. In order to avoid numerical instabilities, RT were rescaled from millisecond to tenths of a second during all numerical analysis. All results are presented in ms.

### Modeling

We aimed to answer three questions with the models considered here. First, we investigated which model (i.e. PROSA, SERIA or SERIA_lr_) explained the experimental data best, and whether all important qualitative features of the data were captured by this model. We did not have a strong hypothesis regarding the parametric distribution of the data and hence, comparisons of parametric distributions were only of secondary interest in our analysis. Second, we investigated whether reduced models that kept certain parameters fixed across trial types were sufficient to model the data. Third, we investigated how the probability of a trial type in a block affected the parameters of the model.

#### Model space

Initially, we considered 15 different models as shown in Table 2. Each model was fitted independently for each subject and condition. Since our experimental design included mixed blocks, we allowed for different parameters in pro- and antisaccade trials, i.e., different increase-rate distributions depending on the *trial type* (TT). Under this hypothesis, the PROSA model had 12 free parameters (6 for each TT), whereas the SERIA model required 4 further parameters *(π_e_* and *π_l_* in each TT). The late race SERIA_lr_ model included 16 parameters for the units (8 for each TT). We did not investigate the case that early reactions could trigger antisaccades but rather fixed the probability of an early antisaccade 1 − *π_e_* to 10^−3^. The rationale behind this was that if early reactions are a priori assumed to never trigger antisaccades, rare but possible early antisaccades might cause large biases when fitting a model.

**Table 2.**
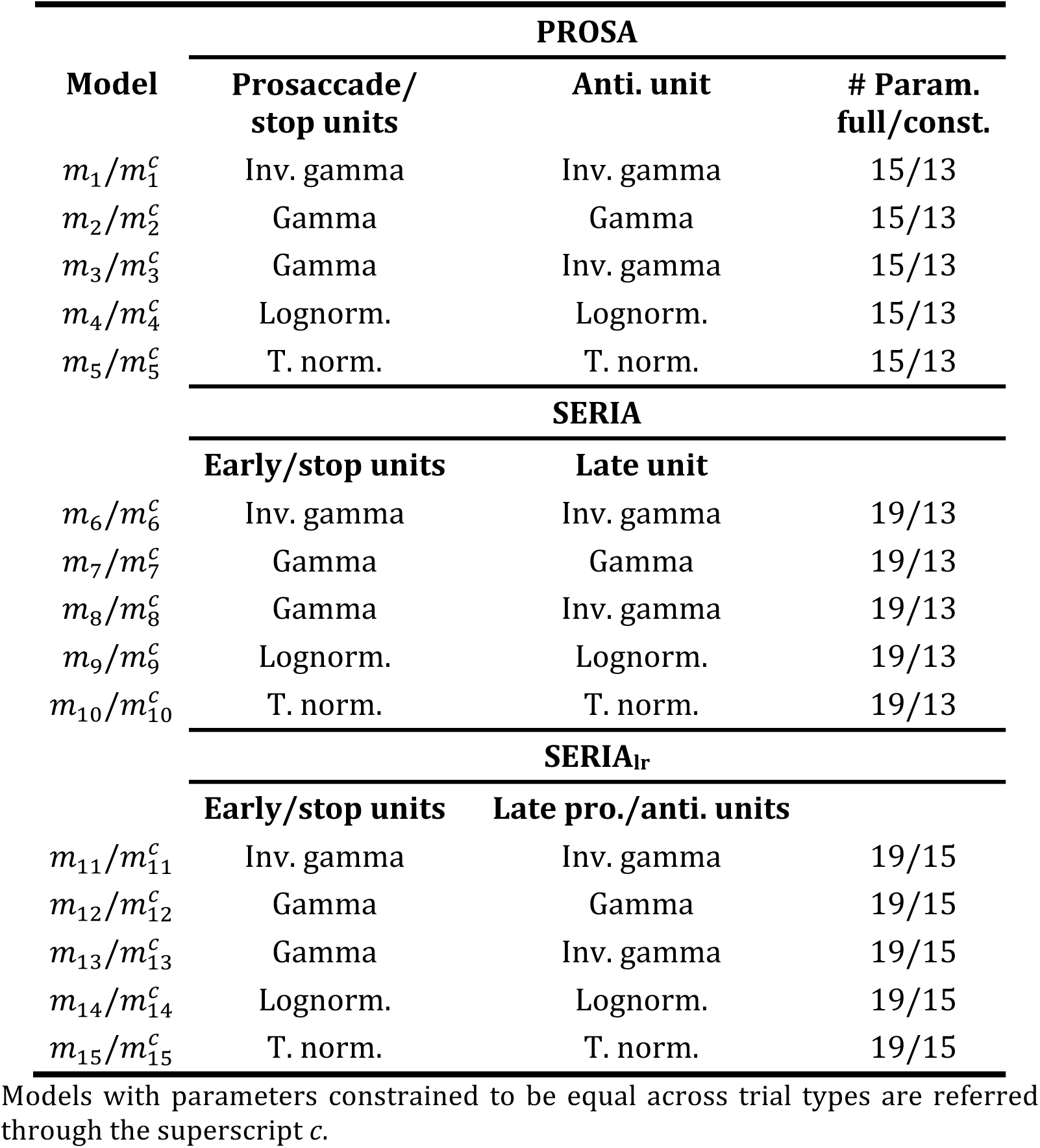
Model families with the respective increase-rate distributions.

| **PROSA** |  |  |  |
| --- | --- | --- | --- |
| **Model** | **Prosaccade/**<br>**stop units** | **Anti. unit** | **# Param.**<br>**full/const.** |
| $m_1/m_1^c$ | Inv. gamma | Inv. gamma | 15/13 |
| $m_2/m_2^c$ | Gamma | Gamma | 15/13 |
| $m_3/m_3^c$ | Gamma | Inv. gamma | 15/13 |
| $m_4/m_4^c$ | Lognorm. | Lognorm. | 15/13 |
| $m_5/m_5^c$ | T. norm. | T. norm. | 15/13 |
| **SERIA** |  |  |  |
|  | **Early/stop units** | **Late unit** |  |
| $m_6/m_6^c$ | Inv. gamma | Inv. gamma | 19/13 |
| $m_7/m_7^c$ | Gamma | Gamma | 19/13 |
| $m_8/m_8^c$ | Gamma | Inv. gamma | 19/13 |
| $m_9/m_9^c$ | Lognorm. | Lognorm. | 19/13 |
| $m_{10}/m_{10}^c$ | T. norm. | T. norm. | 19/13 |
| **SERIA<sub>lr</sub>** |  |  |  |
|  | **Early/stop units** | **Late pro./anti. units** |  |
| $m_{11}/m_{11}^c$ | Inv. gamma | Inv. gamma | 19/15 |
| $m_{12}/m_{12}^c$ | Gamma | Gamma | 19/15 |
| $m_{13}/m_{13}^c$ | Gamma | Inv. gamma | 19/15 |
| $m_{14}/m_{14}^c$ | Lognorm. | Lognorm. | 19/15 |
| $m_{15}/m_{15}^c$ | T. norm. | T. norm. | 19/15 |

Models with parameters constrained to be equal across trial types are referred through the superscript *c*.

Regarding the non-decision time *δ* antisaccade cost *δ_a_*, and rate of outliers *η*, we assumed equal parameters in both TT. Consequently, the full PROSA model had 15 free parameters whereas the full SERIA and SERIA_lr_ models had both 19 free parameters.

In addition to the full models, we evaluated restricted versions of each of them by constraining some parameters to be equal across TT. In the case of the SERIA model, we hypothesized that the parameters of all units were equal, irrespective of TT (i.e., that the rate of the units was not affected by the cue presented in a trial). However, we assumed that the probability that an early or late response was a prosaccade was different in pro- and antisaccade trials. Therefore, in the case of the SERIA model, instead of 12 unit parameters (6 per TT), the restricted model had only 6 parameters for the units’ rates. The parameters *π_e_* and *π_l_* were allowed to differ in pro and antisaccade trials. In the case of the restricted SERIA_lr_ model, the units that underlie the late decision process were allowed to vary across TT, yielding a restricted model with 4 parameters for the early and inhibitory units, and 8 for the late decision process, half of them for each trial type. In the case of the PROSA model, similarly to [17], it is possible to assume that the parameters of the prosaccade unit remain constant across TT, and that the parameters of the stop and antisaccade units depend on TT, yielding 10 parameters for the units.

#### Prior distributions for model parameters

To complete the definition of the models, the prior distribution of the parameters was specified. This distribution reflects beliefs that are independent of the data and provides a form of regularization when inverting a model. In order to avoid any undesired bias regarding the parametric distributions considered here, we reparametrized all but the truncated normal distribution in terms of their mean and variance. We then assumed that the log of the mean and variance of the rate of the units were equally normally distributed (see Table 3). Therefore, the parametric distributions had the same prior in terms of their first two central moments. In the case of the truncated normal distribution, instead of an analytical transformation between its first two moments and its natural parameters *μ* and *σ*^2^, we defined the prior distribution as a density of *μ* and ln *σ*^2^. To ensure that *μ* was positive with high likelihood (96%) we assumed that *μ* ~ *N*(0.55,0.09). The variance term was distributed as displayed in Table 3. As a further constraint, we restricted the parameter space to enforce that the first two moments of the distributions of rates and RTs existed. We relaxed this constraint for the late units of the SERIA_lr_ in order to allow for ‘flat’ distributions with possibly infinite mean and variance. This can describe a case in which the increase rate of one of the late units is extremely low.

**Table 3.** Prior probability density functions.

| Parameter | Probability density function | Expected value | Variance |
| --- | --- | --- | --- |
| $\mu_r$ | $\mathcal{N}(\ln \mu_r; -1.08, 0.97)$ | 0.55 | 0.5 |
| $\sigma_r^2$ | $\mathcal{N}(\ln \sigma_r^2; -2.64, 0.69)$ | 0.1 | 0.01 |
| $\delta$ | $\mathcal{N}(\ln \delta; -1.58, 1.79)$ | 0.5 | 1.25 |
| $\delta_a$ | $\mathcal{N}(\ln \delta_a; -0.87, 1.17)$ | 0.75 | 1.25 |
| $\pi_e$ | $Beta(\pi_e; 0.5, 0.5)$ | 0.5 | 0.145 |
| $\pi_l$ | $Beta(\pi_l; 0.5, 0.5)$ | 0.5 | 0.145 |
| $\eta$ | $Beta(\eta; 0.5, 0.5)$ | 0.5 | 0.145 |

For the non-decision time *δ* and the antisaccade cost *δ_a_*, the prior of their log transform was a normal distribution, consistent across all models. Note that the scale of the parameters *δ* and 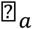 in Table 3 is tenths of a second. The fraction of early outliers *η*, and early and late prosaccades *π_e_* and *π_l_* were assumed to be Beta distributed, with parameters 0.5 and 0.5. Thus, for example, the prior probability of an early outlier is given by

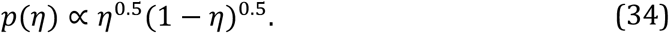

This parametrization constitutes the minimally informative prior distribution, as it is the Jeffrey’s prior *of η, π_e_* and *π_l_*. Table 3 displays the parameters used for the prior distributions.

### Bayesian inference

Inference on the model parameters was performed using the Metropolis-Hastings algorithm [31]. To increase the efficiency of this sampling scheme, we iteratively modified the proposal distribution during an initial ‘burn-in’ phase as proposed by [32]. Moreover, we extended this method by drawing from a set of chains at different temperatures and swapping samples across chains. This method, called population MCMC or parallel tempering, increases the statistical efficiency of the Metropolis-Hasting algorithm [33] and has been used in similar contexts before [34]. We simulated 16 chains with a 5-th order temperature schedule [35]. For all but the models including a truncated normal distribution, we drew a total of 4.1×10^4^ samples per chain, from which the first 1.6×10^4^ samples were discarded as part of the burn-in phase. When a truncated normal distribution was included (models m_5_, m_10_, and m_15_), the total number of samples was increased to 6×10^4^, from which 2×10^4^ were discarded. The convergence of the algorithm was assessed using the Gelman-Rubin criterion [33,36] such that the *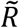* statistic of the parameters of the model was aimed to be below 1.1. When a simulation did not satisfy this criterion, it was repeated until 99.5 percent of all inversions satisfied it.

Models were scored using their log marginal likelihood or log model evidence (LME). This is defined as the log probability of the data given a model after marginalizing out all its parameters. When comparing different models, the LME corresponds to the log posterior probability of a model under a uniform prior on model identity. Thus, for a single subject with data*y*, the posterior probability of model *k*, given models 1 to *n* is

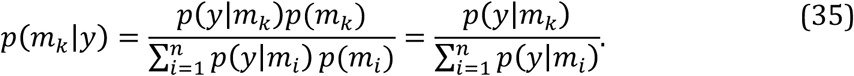

Importantly, this method takes into account not only the accuracy of the model but also its complexity, such that overparameterized models are penalized [37]. A widely used approximation to the LME the Bayesian Information Criterion (BIC) which, although easy to compute, has limitations (for discussion, see [38]). Here, we computed the LME through sampling using thermodynamic integration [33,39]. This method provides robust estimates and can be easily computed using samples obtained through population MCMC.

One important observation here is that the LME is sensitive to the prior distribution, and thus can be strongly influenced by it [40]. We addressed this issue in two ways: On one hand and as mentioned above, we defined the prior distribution of the increase rates of all models in terms of the same mean and variance. This implies that the priors were equal up to their first two moments, and hence all models were similarly calibrated. On the other hand, we complemented our quantitative analysis with qualitative posterior checks [33] as shown in the results section.

Besides comparing the evidence of each model, we also performed a hierarchical or random effects analysis described in [38,41]. This method can be understood as a form of soft clustering in which each subject is assigned to a model using the LME as assignment criterion. Here, we report the expected probability of the model *r_i_*, which represents the percentage of subjects that is assigned to the cluster representing model *i*. This hierarchical approach is robust to population heterogeneity and outliers, and complements reporting the group-level LME. Finally, we compared families of models [42] based on the evidence of each model for each subject summed across conditions.

### Classical statistics

In addition to a Bayesian analysis of the data, we used classical statistics to investigate the effect of our experimental manipulation on behavioral variables (mean RT and ER) and the parameters of the models. We have suggested previously [11,43,44] that generative models can be used to extract hidden features from experimental data that might not be directly captured by, for example, standard linear methods or purely data driven machine learning techniques. In this sense, classical statistical inference can be boosted by extracting interpretable data features through Bayesian techniques.

Frequentist analyses of RT, ER, and parameter estimates were performed using a mixed effects generalized linear model with independent variables *subject* (SUBJECT), *prosaccade probability* (PP) with levels PP20, PP50 and PP80, and when pro- and antisaccade trials were analyzed together, *trial type* (TT). The factor SUBJECT was always entered as a random effect, whereas PP and TT were treated as categorical fixed effects. In the case of ER, we used the probit function as link function.

Analyses were conducted with the function *fitglme.m* in MATLAB 9.0. The significance threshold *α* was set to 0.05.

### Implementation

All likelihood functions were implemented in the *C* programming language using the GSL numerical package (v.1.16). Integrals without an analytical form or well-known approximations were computed through numerical integration using the Gauss-Kronrod- Patterson algorithm [45] implemented in the function *gsl_integration_qng*. The sampling routine was implemented in MATLAB (v. 8.1) and is available as a module of the open source software package TAPAS (www.translationalneuromodeling.org/tapas).

## Results

### Behavior

Forty-seven subjects (age: 23.8 ± 2.9) completed all blocks and were included in further analyses. A total of 27072 trials were recorded, from which 569 trials (2%) were excluded (see Table 4).

**Table 4.** Summary of trials per subject

|  | Valid | Blink | Missing | Aborted | FE | Late S. | Early S. | Total |
| --- | --- | --- | --- | --- | --- | --- | --- | --- |
| Total | 26503 | 188 | 60 | 42 | 249 | 0 | 30 | 27072 |
| Mean | 563.9 | 4.0 | 1.3 | 0.9 | 5.3 | 0.0 | 0.6 | 576 |
| Std. | 9.9 | 5.1 | 2.5 | 1.5 | 5.0 | 0.0 | 1.3 | - |
| Min. | 536 | 0 | 0 | 0 | 0 | 0 | 0 | - |
| Max. | 576 | 22 | 15 | 6 | 19 | 0 | 8 | - |

FE: Fixation errors. Late saccades are saccades elicited after 800ms. Early saccades are prosaccades elicited before 50ms after CP or antisaccades elicited before 110ms after CP.

Both ER and RT showed a strong dependence on PP (Fig 5 and Table 5). Individual data is included in the S1 File and is displayed in S2 Figure. The mean RT of correct pro- and antisaccade trials was analyzed independently with two ANOVA tests with factors SUBJECT and PP. We found that in both pro- (F_2,138_ = 46.9, *p* < 10^−5^) and antisaccade trials (F_2,138_ = 37.3, *p* < 10^−5^) the effect of PP was significant; with higher PP, prosaccade RT decreased, whereas the RT of correct antisaccades increased. On a subject-by-subject basis, we found that between the PP20 and PP80 conditions, 91% of the participants showed increased RT in correct antisaccade trials, while 81% demonstrated the opposite effect (a decrease in RT) in correct prosaccade trials. Similarly, there was a significant effect of PP on ER in both prosaccade (F_2,138_ = 376.1, *p* < 10^−5^) as well as in antisaccade (F_2,138_ = 347.0, *p* < 10^−5^) trials. This effect was present in all but one participant in antisaccade trials and in all subjects in prosaccade trials. Exemplary RT data of one subject in the PP50 condition is displayed in Fig 6.

**Fig 5.**
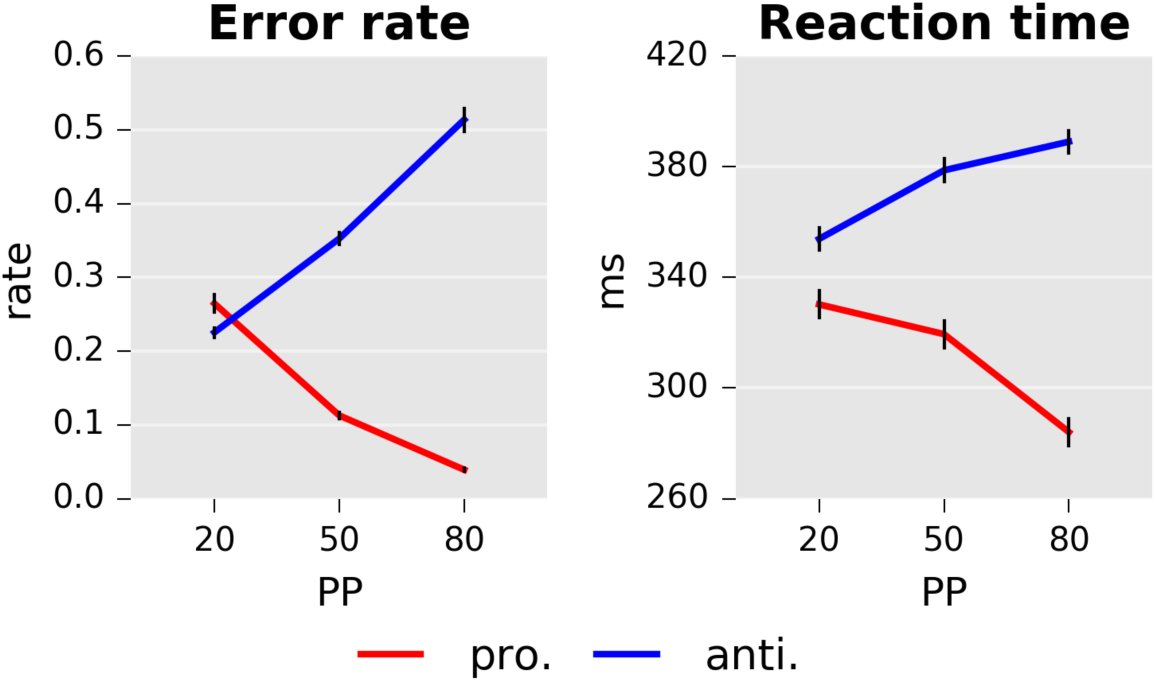
Error rate and reaction times as a function of prosaccade trial probability (PP). Left panel: Mean error rates for pro and antisaccades. Right panel: Mean reaction time in *ms*. Error bars indicate standard errors of the mean. Only correct responses are displayed.

**Table 5.**
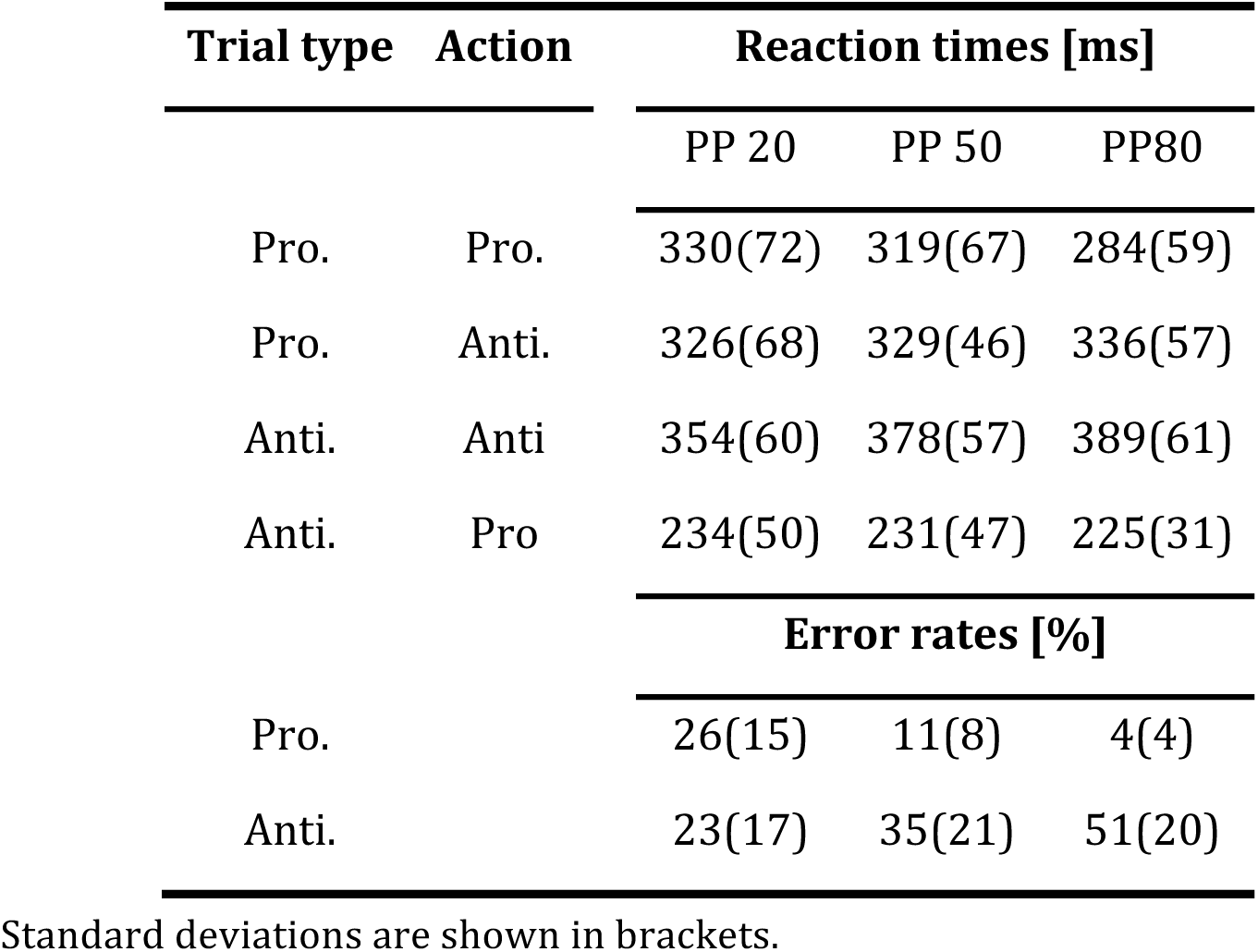
Summary of mean RTs and ERs.

**Fig 6.**
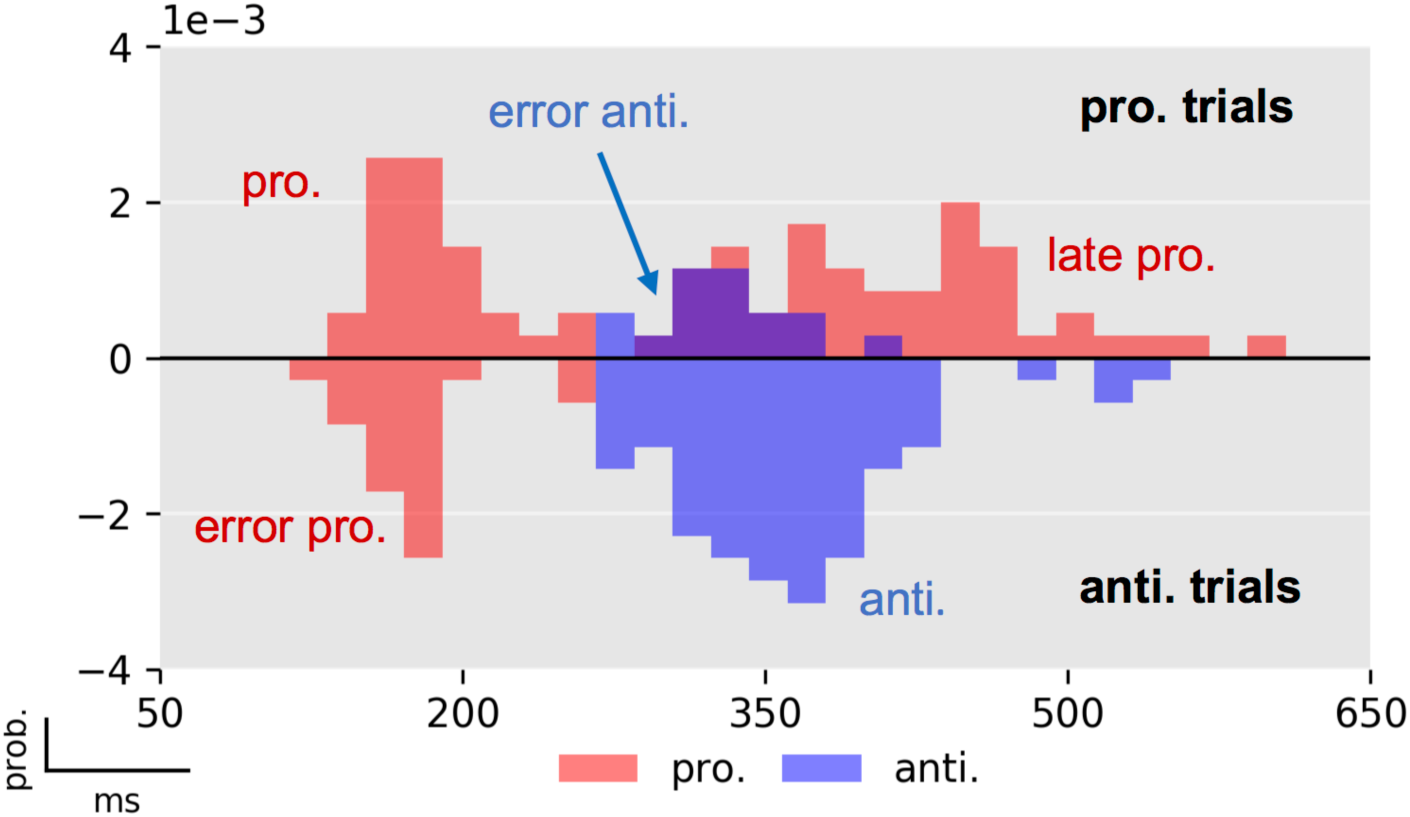
Exemplary RT histogram of one subject in the PP50 condition. Prosaccade trials are displayed in the upper half plane and antisaccade trials in the lower (negative) half plane. Prosaccade actions are depicted in red color, whereas antisaccade actions are shown in blue. Errors in prosaccade trials are antisaccades that for this subject occurred after the first peak of early prosaccades. Errors in antisaccade trials (lower half plane) occurred at a similar latency as early prosaccades in prosaccade trials. The histograms have been normalized to have unit probability mass, i.e., the sum of the area of all bars is one.

### Modeling

#### Model comparison results

Initially, we considered the models outlined in Table 2. The LME over all participants (fixed effects analysis) and the posterior probability of all models and all subjects are presented in Fig 7. Independently of the particular parametric distribution of the units, the SERIAlr models had higher evidence compared to the PROSA and SERIA models. A random effects, family-wise model comparison [42] resulted in an expected frequency of *r* = 87% for the SERIA_lr_ family, *r* = 11% for the SERIA family, and *r*=2% for the PROSA family. In addition, constraining the parameters to be equal across trial types increased the model evidence irrespective of the parametric distribution assigned to the units (Fig 7). Here, the family-wise model comparison showed that models with constrained parameters had an expected frequency of *r* = 98%. Over all 30 models, 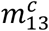 (SERIA_lr_ with constrained parameters, early and inhibitory increase rates gamma distributed, and late units’ rate inverse gamma distributed) showed the highest LME with *ΔLME >* 200 compared to all other models. Following [40], a difference in LME larger than 3 corresponds to strong evidence.

**Fig 7.**
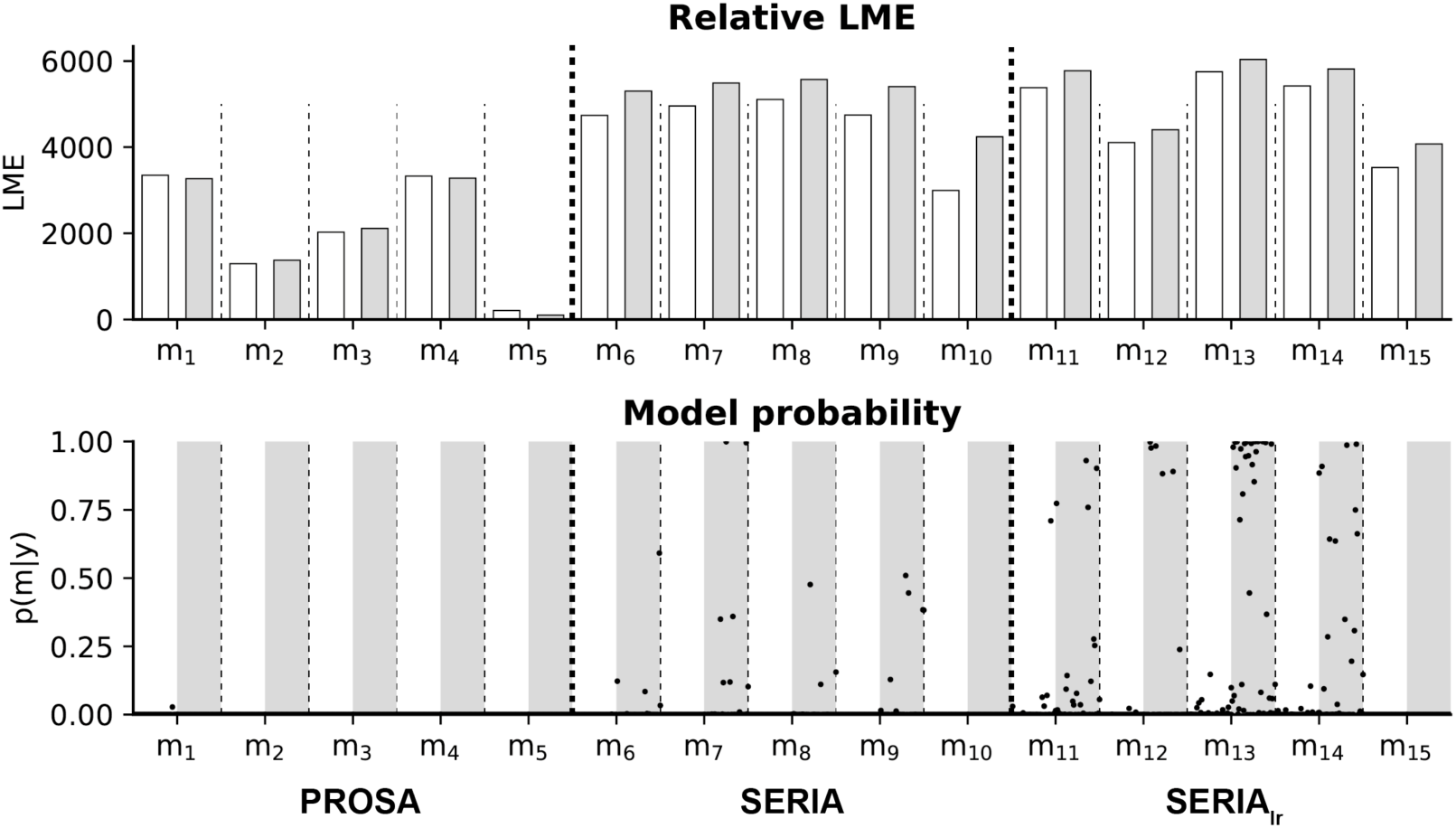
Summary of model comparison. Top: Summed LME of all subjects for all 30 models. White bars show models with all parameters free, grey bars models with constrained parameters. LMEs are normalized by subtracting the lowest LME (m_5_). Model 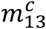 (constrained SERIAlr) exceeded all other models 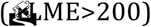. Bottom: Illustration of model probability for all subjects. The posterior model probabilities for all subjects are shown as black dots. In white shading are models with all parameters free, grey bars represent models with restricted parameters. Note that in nearly all subjects, the SERIAlr models with restricted parameters showed high model probabilities.

To verify that the SERIA_lr_ family was not preferred simply because the probability of early prosaccades was fixed, we considered models in the SERIA family with the same property (not displayed). We found that although fixing this value increased the LME of the SERIA family, there was still a difference of *ΔLME >* 90 when comparing the best model of the SERIA_lr_ family and the best model of the SERIA family with a fix probability of early prosaccades.

Fits of four subjects using the posterior samples of the best PROSA (*m*_1_), SERIA 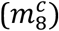, and SERIA_lr_ 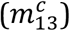 models are depicted in Fig 8. Although model *m*_1_ was the best model in the PROSA family, it clearly did not explain the apparent bimodality of the prosaccade RT distributions. Instead, RTs were explained through wider distributions. No obvious difference could be observed between the SERIA and SERIA_lr_ models. We further examined the model fits in Fig 9 and Fig 10 by plotting the weighted fits and cumulative density functions of the reciprocal RT in the probit scale (reciprobit plot [22]) collapsed across subjects for the best model of each family. The histograms of RTs clearly show a large number of late prosaccades whose distribution is similar to the distribution of antisaccade RTs. The most pronounced - but still small - difference between the SERIA and SERIA_lr_ models was visible in prosaccade trials in the PP20 condition (left panel, upper half plane), in which antisaccade errors displayed lower RT than correct late prosaccades.

**Fig 8.**
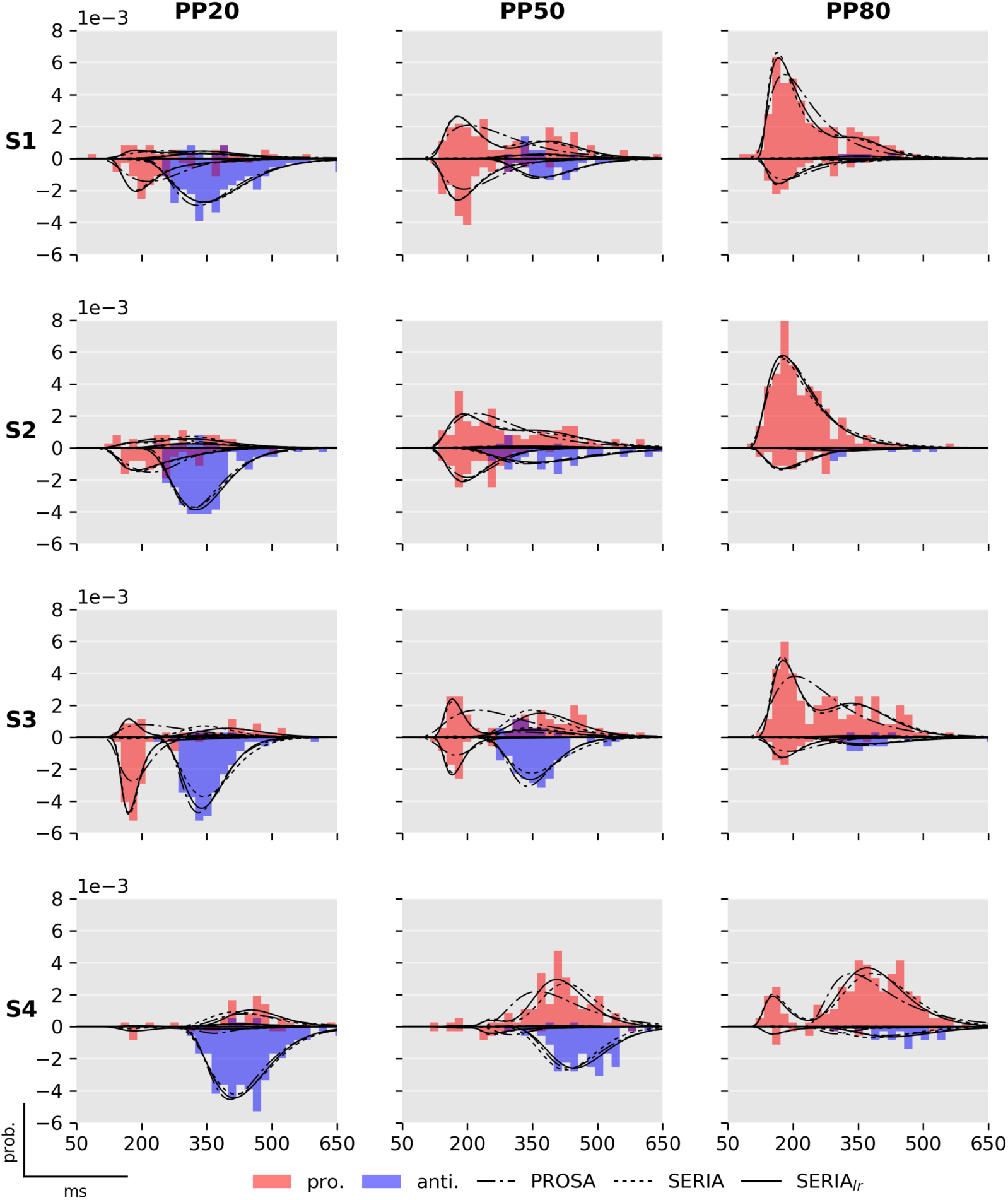
Fits of best PROSA (*m*_1_), SERIA 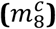 and SERIA_lr_ 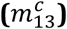 models. Columns display the normalized histogram of the RTs of pro- [red] and antisaccades [blue] in each of the conditions. Rows correspond to individual subjects named S1 to S4 for display purpose. As in Fig. 6, prosaccade trials are displayed on the upper half plane, whereas antisaccade trials are displayed in the lower half plane. The predicted RT distributions based on the samples from the posterior distribution are displayed in solid (SERIA_lr_), broken [SERIA], and dash-dotted [PROSA] lines. Note that data from subject 3 in the PP50 condition is the same as shown in Fig. 6. Early outliers are not displayed.

**Fig 9.**
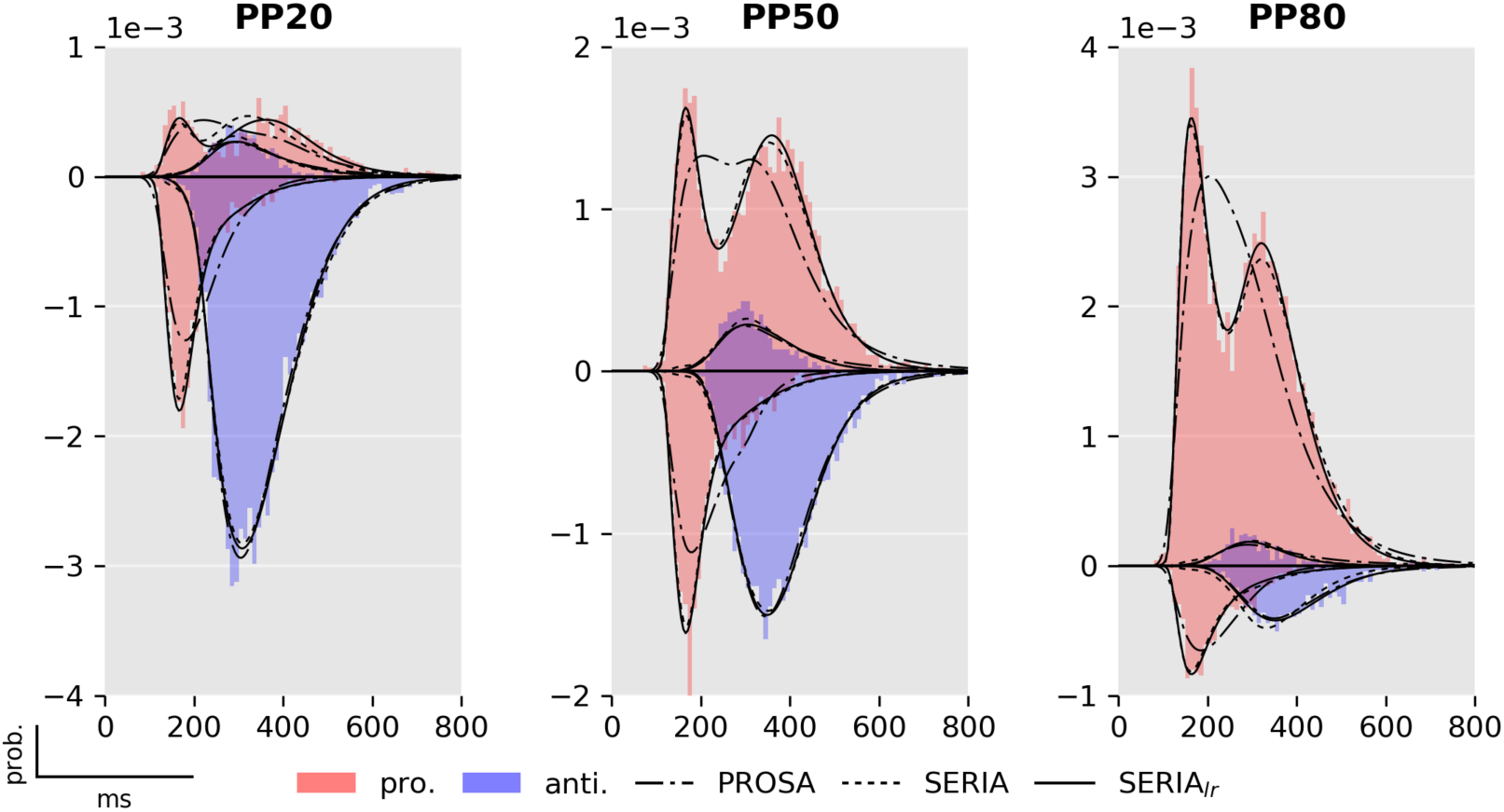
Fits from the best models in each family 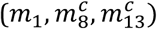. Model fits and RT histograms for each condition collapsed across subjects. For more details see Fig. 6 and 8.

**Fig 10.**
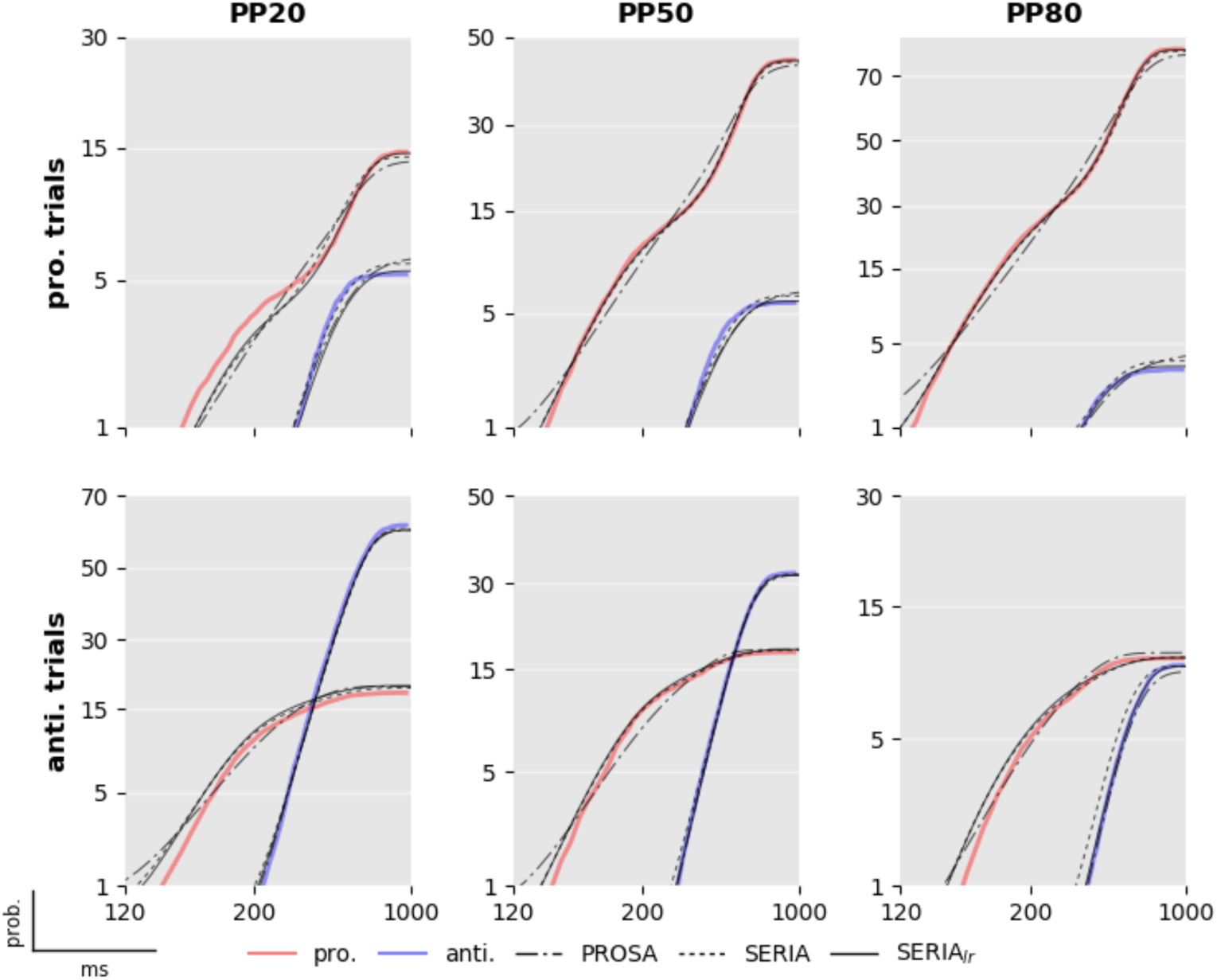
Reciprobit of best models. Predicted and empirical cumulative density function of the reciprocal RT in the probit scale for each condition and model collapsed across all subjects. The data shown are the same as in Fig 9, but split for trial types and illustrated as cumulative distributions. Note that the y-axis is in the probit scale and that nearly all differences between the model and the data occur at very small probability values of 5% or below.

#### Corrective antisaccades

The RTs of antisaccades that follow an error prosaccade were not directly modeled. However, we hypothesized that corrective antisaccades are delayed late antisaccade actions, whose distribution is given by the *response time* distribution of late antisaccades

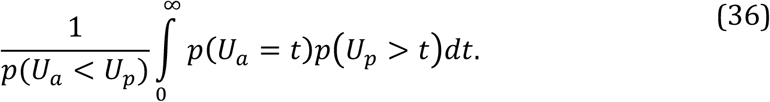

A total of 2989 corrective antisaccades were included in the analysis. The mean (±std) end time of the erroneous prosaccades was 268(±63)ms. The mean RT of corrective antisaccades was 447(±103)ms, and the weighted mean arrival time of the late antisaccade unit was 367ms. Fig 11 displays the histogram of the end time of all prosaccade errors, the RT of all corrective antisaccades and the time shifted (+80ms) predicted response time of late antisaccades. Since we did not have a strong hypothesis regarding the magnitude of the delay of the corrective antisaccades, we selected the time shift to be the difference between the mean corrective antisaccade RT and the mean predicted response time of late antisaccades. Visual inspection strongly suggests that the distribution of corrective antisaccade RTs is well approximated by the distribution of the late responses. The short difference between corrective antisaccades’ RT and the expected response time of the late antisaccade unit (80ms) favors the hypothesis that the plan for a corrective antisaccade is initiated before the incorrect prosaccade had finished.

**Fig 11.**
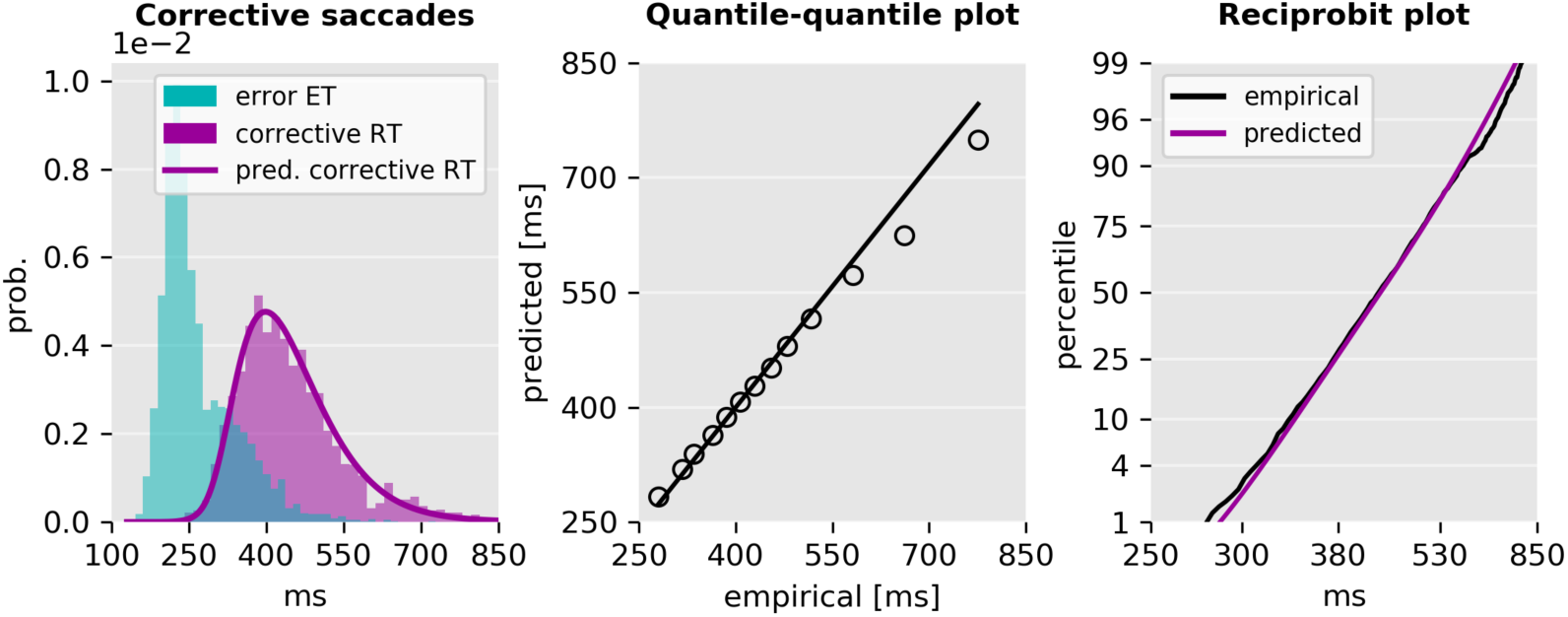
Empirical and predicted RT of corrective antisaccades. Left: End time of erroneous prosaccades, RTs of corrective antisaccades, and time shifted predicted response time distribution of late antisaccades. The time shift was selected to be the difference between the empirical and predicted mean response time. Center: Quantile-quantile plot of the predicted and empirical distribution of corrective antisaccades, and a linear fit to the central 98% quantiles. There is a small deviation only at the tail of the distribution. Right: Reciprobit plot of the empirical and predicted cumulative density functions of the RT of corrective antisaccades. The scale of the horizontal axis is proportional to the reciprocal RT. The vertical axis is in the probit scale.

#### Effects of prosaccade probability on model parameters

The effect of PP on the parameters of the model was investigated by examining the expected value of the parameters of the best scoring model *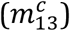*. Initially, we considered the question of whether the mean arrival or response time of each of the units changed as a function of PP. For arrival times, this corresponds to

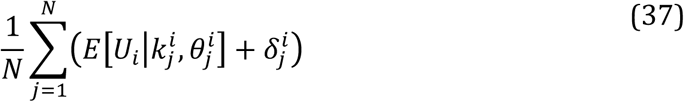

where *i* is an index over the units, *j* is an index over *N* samples collected using MCMC, and 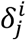 is the estimated delay. In the case of the late units, we considered only the response time of correct actions. Fig 12 left displays the mean arrival and response times. These were submitted to four separate ANOVA tests, which revealed that PP had a significant effect on all four units: early unit *(F*_2,138_ = 9.2, *p* < 10^−3^), late antisaccade (*F*_2,138_ = 26.6, *p* < 10^−3^), late prosaccade (*F*_2,138_ = 19.6, *p* < 10^−3^), and inhibitory unit (*F*_2,138_ = 30.9, *p* < 10^−3^). We then explored the differences across conditions through planned post hoc tests on each condition for each of the units (Table 6). The arrival times of the early unit did not change significantly between condition PP20 and PP50, but decreased significantly in the PP80 condition as compared to the PP50 block. The response times of late antisaccades increased significantly between the PP20 and the PP50 conditions but not so between the PP50 and PP80. Late prosaccades followed the opposite pattern, showing only a significant decrease in response time between the PP50 and PP80 conditions. Finally, the inhibitory unit changed significantly across all conditions.

**Fig 12.**
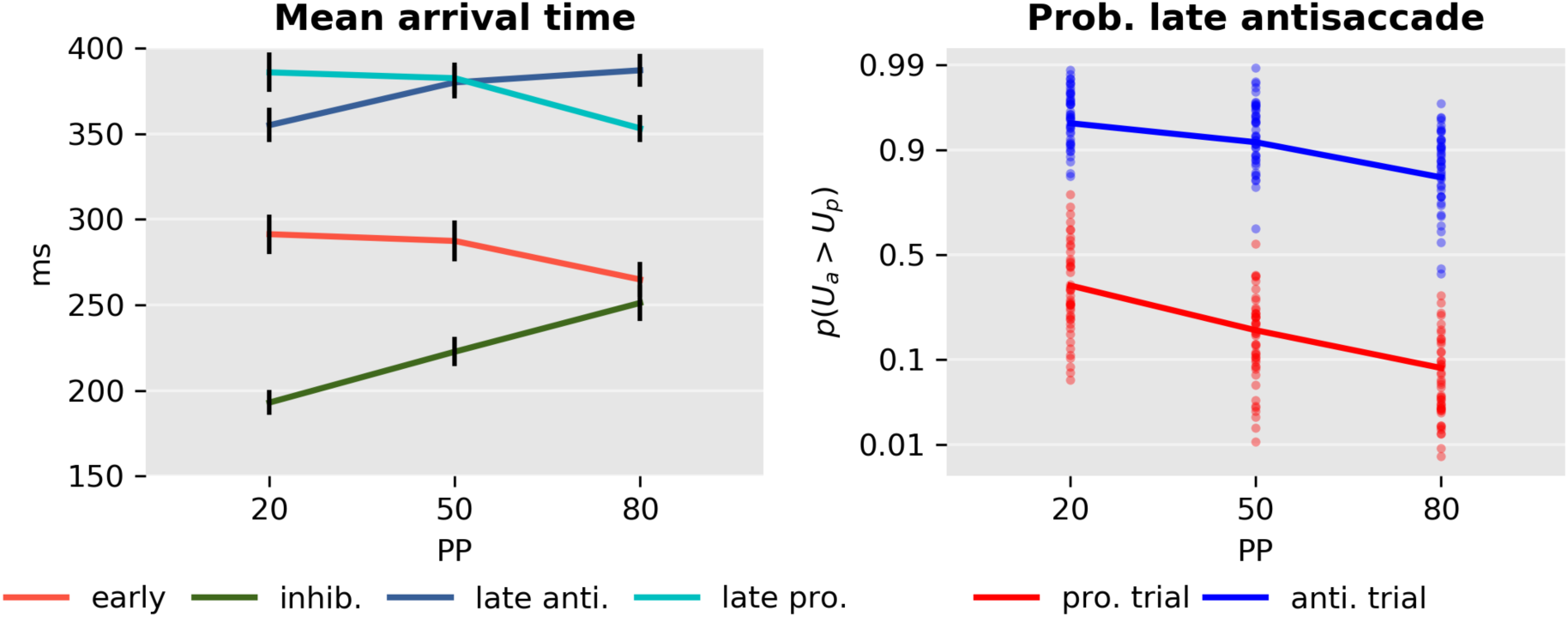
Model parameters. Left: Mean arrival or response time and standard error of the early and inhibition units and late pro- and antisaccades. Right: Probability of a late antisaccade *p*(*U_a_* > *U_p_*) in prosaccade (red) and antisaccade (blue) trials in each condition in the probit scale.

**Table 6.** Post hoc comparison of the effect of PP.

|  | Early |  |  | Inhib. |  |  | Late pro. |  |  | Late anti. |  |  |
| --- | --- | --- | --- | --- | --- | --- | --- | --- | --- | --- | --- | --- |
| | Mean | $t_{138}$ | p | Mean | $t_{138}$ | p | Mean | $t_{138}$ | p | Mean | $t_{138}$ | p |
|  | [ms] |  |  | [ms] |  |  | [ms] |  |  | [ms] |  |  |
| PP20-PP50 | 4 | 0.6 | 0.50 | -30 | -4.0 | <0.001* | 3 | 0.5 | 0.55 | -24 | -36 | <0.001* |
| PP20-PP80 | 26 | 3.9 | <0.001* | -58 | -7.8 | <0.001* | 32 | 5.7 | <0.001* | -32 | -5.4 | <0.001* |
| PP50-PP80 | 22 | 3.3 | 0.001* | -28 | -3.8 | <0.001* | 29 | 5.1 | <0.001* | -7 | -1.5 | 0.13 |

Effect of PP on the mean arrival time for each of the early, inhibitory units, and the late pro and antisaccade units in the corresponding trial type.

Finally, we examined how the probability of a late antisaccade *p*(*U_a_* > *U_p_*) (Fig 12, right) depended on PP and TT. The estimated parameters for both pro- and antisaccade trials were analyzed with a model with factors SUBJECT, TT, PP and the interaction between TT and PP. An ANOVA test demonstrated that both PP (*F*_2,276_ = 51.2, *p* < 10^−3^) and TT (*F*_1,276_ = 985.0, *p* < 10^−3^) had a significant effect, but there was no evidence for an interaction between the two factors (*F*_2,276_ = 1.5, *p* = 0.23), suggesting that PP affected the probability of a late antisaccade equally in pro- and antisaccade trials.

### Subject specific parameters

Finally, we investigated how some of the parameters of the model were related to each other across subjects. Because it has been commonly reported that schizophrenia is related with higher ER, but also with increased antisaccade RT, an interesting question is whether higher late-action response times are correlated with the percentage of late errors and inhibition failures, i.e., early saccades that are not stopped. We found that the response time of late pro (*F*_1,135_ = 13.6, *p* < 0.001) and antisaccades (*F*_1,135_ = 7.1, *p* < 0.01) was negatively correlated with the probability of a late error (Fig 13), but no significant interaction between PP and response time was found (pro: *F*_2,135_ = 1.7, *p* = *p* = 0.19; anti *F*_2,135_ = 0.3, *p* = 0.76). Hence, late responders tended to make fewer late errors, suggesting a speed/accuracy trade-off in addition to the main effect of PP. We further considered the question whether the percentage of inhibition failures was correlated with the expected arrival time of the late antisaccade unit in antisaccade trials (Fig. 13 right). Note that the number of inhibition failures is the same in both trial types in a constrained model, but inhibition failures are errors in antisaccade trials and correct early reactions in prosaccade trials. We found that these parameters were not significantly correlated (*F*_2,135_ = 1.2, *p* = 0.26). This was also the case when we considered the expected response time of late prosaccades in prosaccade trials (not displayed; *F*_2,135_ = 0.0, *p* = 0.98).

**Fig 13.**
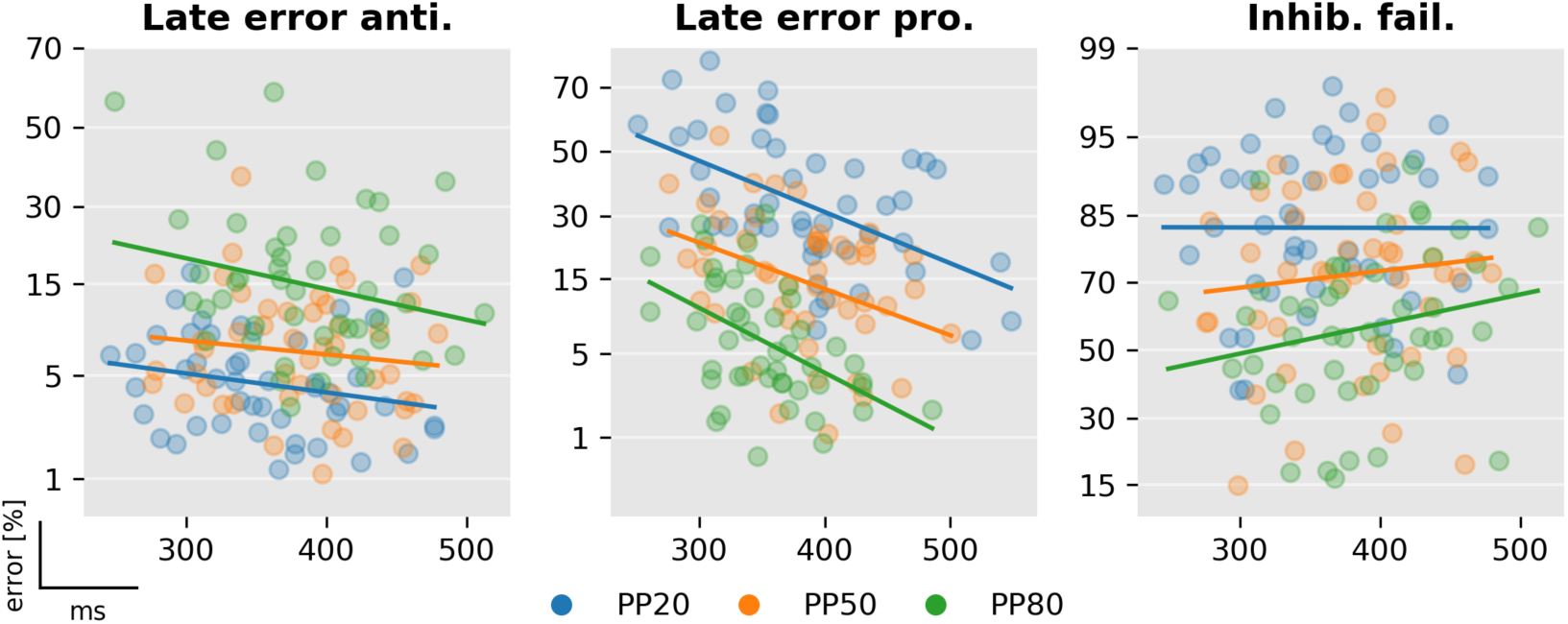
Correlation between late arrival times and errors. Left: Percentage of late errors against late antisaccades’ response times in antisaccade trials. Center: Percentage of late errors against late prosaccades’ response time in prosaccade trials. Left: Percentage of inhibitory failures against late antisaccades’ response time in antisaccade trials. The horizontal axis is in the probit scale.

Fig 14 illustrates the posterior distribution of late errors and inhibition failures of two representative subjects as estimated using MCMC. Clearly, PP induced strong differences in the percentage of inhibition failures and late errors in prosaccade trials in both subjects. The effect of PP is less pronounced in late errors in antisaccade trials. The posterior distributions also illustrate how the SERIA_lr_ model can capture individual differences: For example, the percentage of late prosaccade errors in the PP80 condition and the percentage of inhibition failures across all conditions are clearly different in each subject.

**Fig 14.**
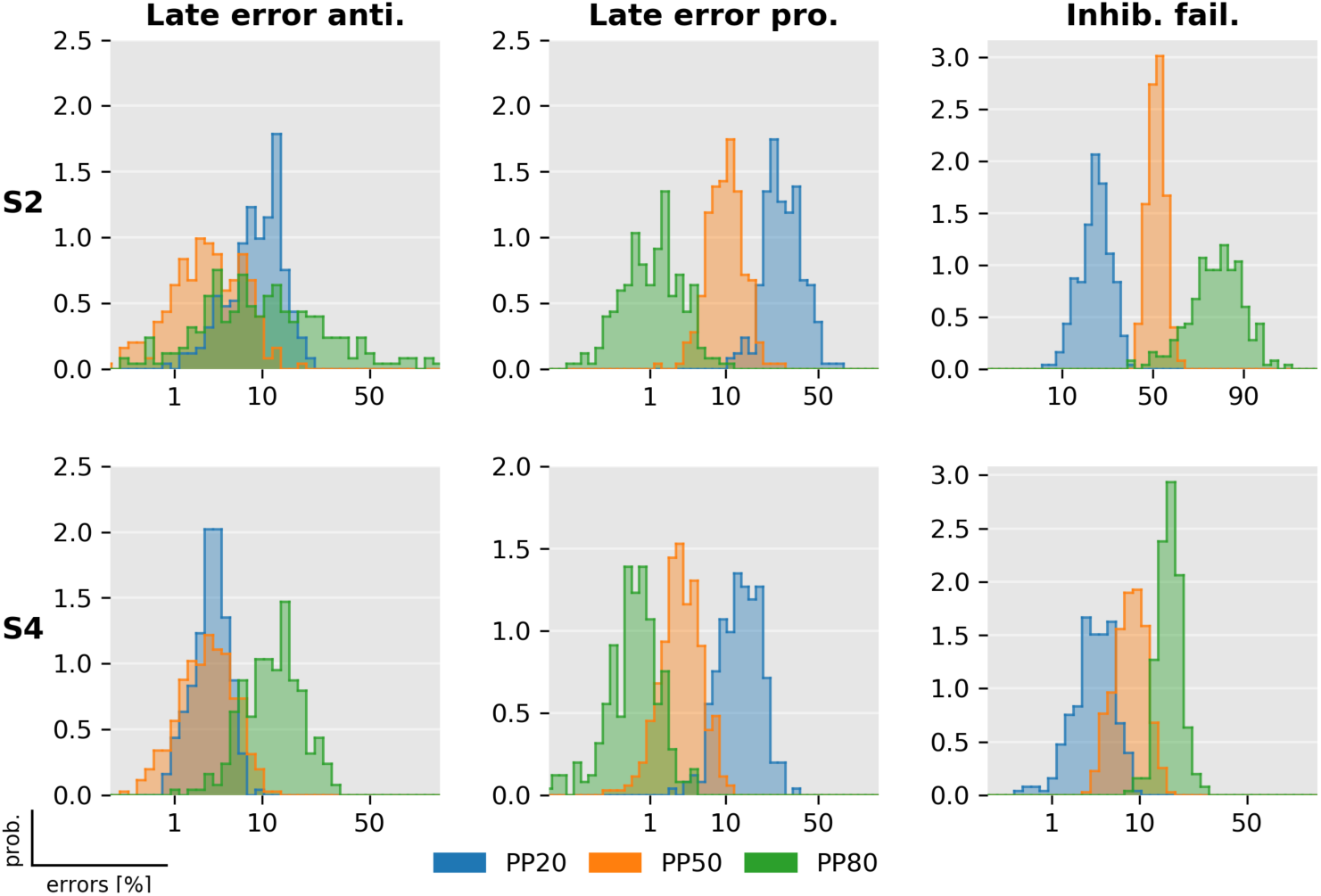
Posterior distribution of late errors and inhibition failures. The posterior distribution of the percentage of late and inhibitory failures of two exemplary subjects (see Fig 8 and 13). Samples from the posterior distribution were obtained using MCMC. Histograms display the distributions of the samples in in the probit scale (horizontal axis). For these two subjects, the posterior distribution of late prosaccade and inhibitory failures clearly discriminates between the three PP conditions.

## Discussion

In this study, we provided a formal treatment of error rates (ER) and reaction times (RT) in the antisaccade task using probabilistic models. We applied these models to data from an experiment consisting of 3 mixed blocks with different probabilities of pro- and antisaccades trials. Model comparison showed that a novel model that allows for late pro- and antisaccades explains our experimental findings better than a model in which all late responses are assumed to be antisaccades. The parameter estimates of the hidden units of the model showed that changes in the inhibition unit and a late decision process explained most of the overt changes in behavior caused by our experimental manipulation, i.e., differences in trial type probability. Moreover, we found that all units were sensitive to the PP in a block, although late responses tended to plateau when the corresponding trial type was not highly frequent.

Our main finding is that two decision processes are necessary to properly model the antisaccade task: on one hand, an early race between a prepotent response towards a target and an endogenously generated signal to cancel this action, and, on the other hand, a secondary late race between two units encoding the cue-action mapping. Although the late decision process can be closely approximated by assuming that RT and actions are independent (at least in our experimental design), Bayesian model comparison demonstrated that late decisions are more accurately described by a race between two units representing different actions. The two decision processes are the sources of early errors-fast prosaccades in antisaccade trials- and late errors-late actions incongruent with the cue presented. The late decision process displays a speed/accuracy tradeoff and is biased by the probability of a trial type in a block. Moreover, this decision process predicts the RT distribution of corrective antisaccades that follow early errors. Because the extra latency of these corrective antisaccades (80ms) is much lower than the average time by which the erroneous prosaccade is terminated, it is unlikely that corrective antisaccades are due to a restart in the decision process, but rather these are late actions that overwrite early errors.

### Influence of trial type probability on reaction times and error rates

Our results show that both RT and ER depend on PP. While this was a highly significant factor in our study, there are mixed findings in previous reports. ER in antisaccade trials was found to be correlated with TT probability in several studies [29,46,47]. However, this effect might depend on the exact implementation of the task [47,48]. Changes in prosaccade ER similar to our study have been reported by [29] and [48]. Studies in which the type of saccade was signaled at fixation prior to the presentation of the peripheral cue do not always show this effect [47]. The results on RTs are less consistent in the literature. Our findings of increased anti- and decreased prosaccade RTs with higher PP are in line with the overall trend in [29], and with studies in which the cue was presented centrally [29,47]. Often, there is an additional increase in RT in the PP50 condition [29,47], which was visible in our data as a slight increase in RT in the PP50 condition on top of the linear effect of PP. Overall, RTs in our study were relatively slow compared to studies in which the TT cue was separated from the spatial cue [46,47]. However, a study with a similar design and added visual search reported even slower RTs in both pro- and antisaccades [29].

### Interpretation of model comparison results

Formal comparison of generative models can offer insight into the mechanisms underlying eye movement behavior [11] and might be relevant in translational neuromodeling applications, such as computational psychiatry [49-53]. Here, we have presented what is, to our knowledge, the first formal statistical comparison of models of the antisaccade task. For this, we formalized the model introduced in [17] and proceeded to develop a novel model that relaxes the one-to-one association of early and late responses with pro- and antisaccades, respectively. All models and estimation techniques presented here are openly available under the GPLv3.0 license as part of the open source package TAPAS (www.translationalneuromodeling.org/tapas).

Bayesian model comparison yielded four conclusions at the family level. First, the SERIA models were clearly favored when compared to the PROSA models. Second, including a late race between actions representing late pro- and antisaccades (SERIA_lr_) resulted in an increase in model evidence, compared to a model not including a late race (SERIA). Third, models in which the race parameters of the early and inhibitory unit were constrained to be equal across TT had a higher LME than models in which all parameters were free. Hence, the effect of the cue in a single trial was limited to the late action, and did not affect the race between an early and inhibitory process. This constitutes an important external validation, as it means that model comparison does favor a model which respects the temporal order of the experiment: Information about TT is only available after the stimulus was presented and, thus, it is unlikely to have an impact of the fast reactive responses. Fourth, early responses were nearly always prosaccades. Crucially, these four conclusions are based on family-wise comparison across all parametric distribution of the increase rate of the units.

A further consequence of our findings is that two independent and qualitatively different decision processes lead to an antisaccade: the race process between early and inhibitory units, and the secondary decision process that generates late responses. A separation of decisions into a ‘where’ and a ‘when’ relationship is not linear, for any continuous in conceptual terms. However, model comparison showed that these two components (‘where’ and ‘when’) cannot be completely dissociated and that time plays a role in the late decision. Nevertheless, the assumption that action type and arrival time of late responses were independent yielded a good fit to this particular data set, suggesting that it is, in many cases, an acceptable approximation to assume a time-independent late decision process. The most obvious difference between the SERIA and SERIA_lr_ can be observed in prosaccade trials in the PP20 condition (left panel, upper half plane Fig. 9), in which late prosaccades are slower than antisaccades. We discuss this point in more detail later.

#### Parametric distribution of reaction times

The parametric distribution of oculomotor RTs has been discussed in great detail in the literature (e.g., [13,55]). Here, we did not aim at determining the most suitable distribution, but rather opted for a practical approach by evaluating different models with a reduced number of parametric distributions. We then based our conclusions on the model with the highest LME. Nevertheless, one can consider the connection of the models presented here with other families of parametric distributions. In particular, the linear relationship

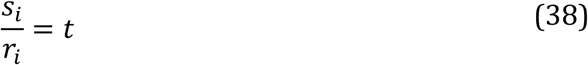

could be seen as to be formally inconsistent with the observation that RT are likely to be explained by stochastic accumulation processes (see for example [56,57], but [58]). This is a weaker constraint than one would expect, because under low noise conditions, for example, the linear relationship can be a good approximation of neural activity. Even if the relationship is not linear, for any continuous function *ϕ* with an inverse function *ϕ* ^−1^, the model can be recasted as [59]:

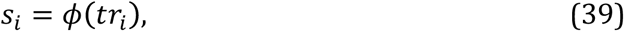

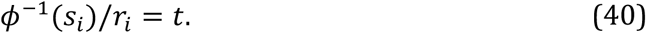

In any case, linear accumulation models have been shown to yield similar conclusions to stochastic accumulation models [58].

More generally, it can be shown that if RTs follow a generalized inverse normal distribution (GIN) of the form

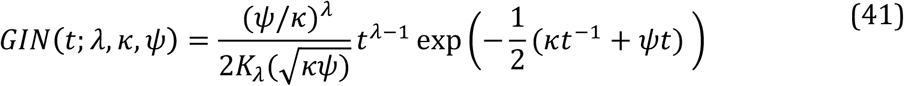

where *λ* ≤ 0, and *K_λ_* is a modified Bessel function of the second kind, there exists a continuous diffusion process whose first hit distribution (FHD) follows the GIN [60]. A particular case of this distribution is the Wald distribution for which 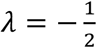. It is the FHD of the Brownian diffusion process with drift

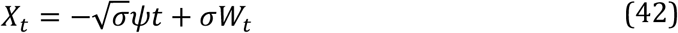

where *W_t_* denotes a Wiener process, *x*_0_ > 0, and the absorbing boundary *a* is zero. More relevant here, when *ψ* = 0 the distribution reduces to an inverse gamma distribution, the FHD of the process

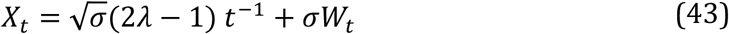

with *x*_0_ > 0 and boundary *a* = 0 (for a detailed mathematical treatment see [60]). Thus, if the rates of a ballistic, linear processes are assumed to be gamma distributed, the RTs follow a distribution that is formally equivalent to a first hit model with stochastic updates and fixed rates. While the model presented here can be seen as a ballistic accumulation model, this equivalence suggests that it is *compatible* with a diffusion process with infinitesimal mean change proportional to *t*^−1^.

#### Other antisaccade models

In broad terms, three families of antisaccade models can be distinguished (reviewed in [61]),The first set of models is based on a race process with independent saccadic and stop units. These models build on the seminal work of [16] on the stop-signal paradigm. According to this model, a ‘GO’ signal triggers a stochastic ‘race’ process that generates a response once it reaches threshold. Critically, a stop signal triggers a second process that inhibits the first ‘GO’ response if it is the first to reach threshold. Importantly, the pace of both units is mutually independent. This model was further extended for the antisaccade task by [17] (but see [14,21], and the review in [20]), who included a third unit such that an antisaccade is generated when a reflexive prosaccade is inhibited by an endogenously-triggered stop process. Note that the original ‘horse-race’ model has also been modified [62] to account for different competing response actions, similarly as in the antisaccade task. The models proposed here belong to this family.

A second type of model relies on lateral or mutual inhibition of competing pro- and antisaccade units. In this direction, Cutsuridis and colleagues [61,63,64] proposed that lateral inhibition is implemented by inhibitory connections in the intermediate layers of the superior colliculus. Thus, saccades are the result of accumulation processes, but these are not independent of each other. Crucially, no veto-like stop signal is required. Although no formal model-fitting has been proposed for this model, qualitative agreement with data suggests that it might capture behavioral patterns relevant in translational applications [64,65]. Since no probabilistic version of this model is available, it is not yet possible to decide on the grounds of model comparison whether mutually dependent or independent race processes best explain current behavioral findings.

Finally, several models that incorporate detailed physiological mechanisms have been proposed [23,66-68]. These models cannot be easily assigned to one of the above categories, as they often employ both an inhibitory mechanism that stops or withholds the reactive responses as well as competition between actions. In addition, while more realistic models possess a more fine-grained representation of the underlying neurobiology, they rely on a large number of parameters and for this reason, it is difficult to fit them to behavioral data (for discussion, see [11]).

Regarding neurobiologically realistic models, the model proposed by [23] is the most similar to the SERIA model. It posits two different mechanisms that interact in the generation of antisaccades: an action-selection module and a remapping module that controls the cue-action mapping. As a consequence, this model allows for the generation of late errors that follow a similar RT distribution as correct antisaccades. Consistent with this observation, the SERIA model can quantitatively distinguish between inhibition and late cue-action mapping errors (Fig 15, left panel). A less obvious similarity between the SERIA model and [23] is that different cues do not lead *directly* to different dynamics in the action module, but only in the so-called ‘remapping’ module. Furthermore, the incorporation of a late race is conceptually close to the approach proposed by [23], which includes a winner take all competition in what we have referred here as late responses. Similarly, our model comparison results show that different cues (i.e., trial types) do not affect the GO/NO-GO process but only the late cue-action mapping.

**Fig 15.**
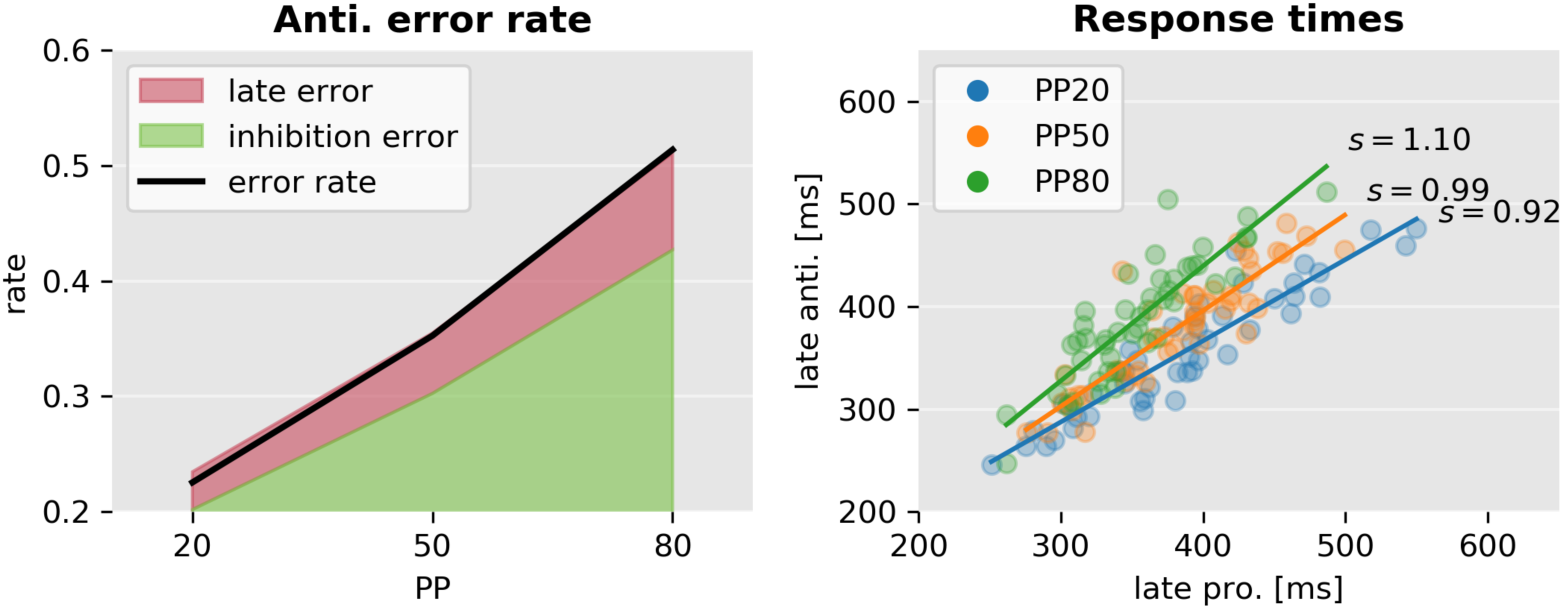
Error sources and the correlation of response times. Left: Error rate (black line) split into the two causes predicted by the model. Inhibition errors are early actions that always trigger prosaccades. Similarly as described by [23], late errors occur when a late response leads to a prosaccade. Right: Correlation between correct antisaccade and late prosaccades response times according to the best SERIA_lc_ model. The best linear fit is depicted as a solid line. The mean ratio of pro- and antisaccade response times (*s*) is displayed on the right. Although late pro - and antisaccade response times are highly correlated, their ratio is different in each condition (interaction PP and late prosaccade response time *F* = 9.2, *p* < 0.001).

### Parameter changes across trial types

One of the most salient results presented here is that models in which the parameters of the units were constrained to be equal across trial types had a larger LME than models in which all the parameters were free, suggesting that the early and inhibitory race units were not affected by the cue presented on a single trial. While visual inspection of the predicted likelihood under the posterior parameters showed that most of the prominent characteristics of the data were explained correctly, some more subtle effects were not captured accurately by the SERIA model. This is particularly clear in the PP20 condition, in which the SERIA model displays a large bias in prosaccades trials in the PP20 condition. One possible explanation is that restricting the parameters across trial types made the model too rigid to capture this effect. Fig 16 compares the fitted RT distributions for models *m*_8_ (SERIA) and *m*_13_ (SERIA_lr_), in which no constraint on the parameters was imposed. Both models are qualitatively almost identical, although as shown in Fig 7, the LME favored the SERIAlr model. Thereby, the qualitative similarity between both models indicates that, in our experiment, the RT of late decisions is only weakly dependent on time. In conclusion, although removing the constraint on the parameters did improve the fit, the differences are marginal and, thus did not justify the additional model complexity. As mentioned above this is consistent with the notion that the information about trial type is only available to the subject once the peripheral stimulus (green bar) has been processed. Presumably tens of milliseconds after the stimulus onset. In fact, this example illustrates the protection against overfitting provided by the LME, as this is a case in which simpler models were preferred over more complex models despite of slightly less accurate fits.

**Fig 16:**
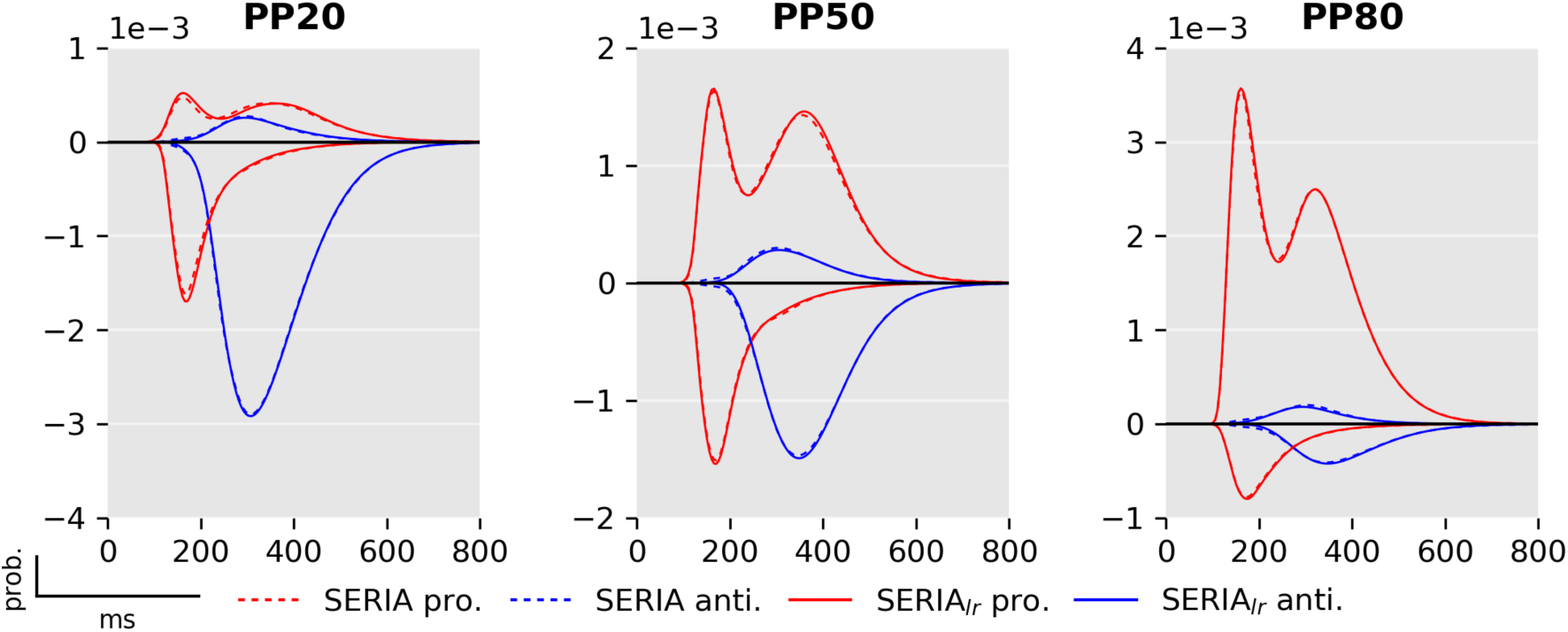
Comparison between unconstrained SERIA and SERIA_lr_ models. Comparison between models *m*_8_ (broken lines; SERIA model) and *m*_13_ (solid lines; late race SERIA_lr_ model.).

Arguably, the constrained SERIA model fails to fully capture the RT of late prosaccade in the PP20 and PP50 conditions because of the assumption that late prosaccades have the same arrival time as late antisaccades. As shown in Fig 15, although the response time of late pro- and antisaccades are strongly correlated, the average ratio of the response times changes across conditions.

### The effect of trial type probability

It is far from obvious why TT probability affects RT and ER in the antisaccade task. One possible explanation is that increased probability leads to higher preparedness for either pro- or antisaccades. Such a theory posits an intrinsic trade-off between preparations for one of the two action types that leads to higher RTs and ERs in low probability trials. Thus, a trade-off theory predicts that the arrival times of early and late responses should be negatively correlated. Although this hypothesis can explain our behavioral findings in terms of summary statistics, our model suggests a more complicated picture.

The main explanation of our results is the effect of TT probability on the inhibitory unit and the probability of a late prosaccade. A higher probability of antisaccade trials leads to faster inhibition and to a higher number of late prosaccades. This resulted in higher mean RT in prosaccade trials when PP is low. In the case of antisaccades, although the mean arrival times of the late unit increased in the PP50 condition, the increased arrival time of the inhibitory unit on the PP80 condition skewed the antisaccade distribution towards higher RTs. Nevertheless, the SERIA_lr_ implies the anticorrelation of late pro- and antisaccades in a single trial type, as these are the results of a GO-GO race.

### Action inhibition

The biological implementation of action inhibition in the antisaccade and other countermanding tasks has received a lot of attention and is still debated [69-73]. Our work adds evidence to the theory that the antisaccade task requires a process that inhibits prepotent responses and is independent of the initiation of a late action [20]. Recent evidence from electrophysiological recordings in the rat brain ([74] reviewed by [71]) suggests that the hypothesized race between go and inhibitory responses might be implemented by different pathways in the basal ganglia [68]. In addition to the basal ganglia, microstimulation of the supplementary eye fields tends to facilitate inhibition of saccades in the countermanding task [75].

### Corrective antisaccades

Although not a primary goal of our model, we considered the question of predicting corrective antisaccades. This problem has received some attention recently [18,61,65,76], as more sophisticated models of the antisaccade task have been developed. We speculated that corrective antisaccades are generated by the same mechanism as late responses. Thus, their RT distribution should follow a similar distribution. Our results strongly suggest that this is the case (see Fig 11). Moreover, the time delay of the corrective antisaccades indicates that, on average, these actions are not the result of the late unit being restarted at the end time of the erroneous prosaccade, as this would lead to much higher RTs. Rather, the planning of a corrective antisaccade might be started much before the end of the execution of an erroneous prosaccade, in accordance with the parallel planning model of the antisaccade task [46] and the ‘GO - STOP+GO’ model in [21].

### Translational applications

Despite the large number of studies of clinical patients using the antisaccade task, an important question remains open: What are the causes of the errors in different neurological and psychiatric conditions? For example [77,78] argued that errors in schizophrenia might be explained, at least partially, by a failure to generate a secondary late action based on several modifications of the antisaccade task. However, it was also proposed that the increased ER in schizophrenia is due to high tonic dopamine levels in the basal ganglia, that lead to decreased inhibition of early responses [68]. More generally, different neurological and psychiatric diseases, or even patients with the same condition, might be characterized by a different source of errors. For example, there is intriguing evidence [79] that patients with different diseases such as attention deficits disorders [80], Parkinons’ disease [81], and amyotrophic lateral sclerosis [82] might be characterized by different ratios of early and late errors. An interesting experimental finding in our study related to this is the considerable amount of erroneous antisaccades in prosaccade trials. An increased number of such errors could be caused by reduced cognitive flexibility leading to impaired shifting between tasks as observed for example in obsessive compulsive disorder [83]. The ability to quantify different types of errors through computational modeling might help to further characterize these diseases.

## Summary

Here we have presented a novel model of the antisaccade task. While the basic structure of the model follows the layout of a previous model [17], we have introduced two crucial advancements. First, we postulated that late responses could trigger both pro- and antisaccades, which are selected by an independent decision process. Second, the generative nature of our model allows for Bayesian model inversion, which enables the comparison of different models and families of models on formal grounds. To our knowledge this has not been done for any of the previous models of the antisaccade task, which is of relevance for translational applications that aim at better understanding psychiatric diseases by means of computational modeling.

The application of the model to a large data set yielded several novel results. First, the early and inhibitory race processes triggered by different cues are almost identical. Moreover, different PP had very different effects on the individual units, which was not obvious from the linear analysis of the mean RT and ER.-Crucially, our modeling approach allowed us to look at a mechanistic explanation or the effects of PP by examining the individual units. In future work we aim to disentangle the mechanisms of behavioral differences caused by neuromodulatory drugs and psychiatric illnesses using formal Bayesian inference.

## Acknowledgments

We thank Saee Paliwal for her remarks, and Jeffrey D. Schall and an anonymous reviewer for their helpful comments on an earlier version of this manuscript.

## Supporting Information

**Supporting information 1**

S1_Dataset.zip. Table of Data. Spreadsheet including all reaction times, actions and errors that entered the analysis. More details are included directly in the file.

## Supporting information 2

**S2:**
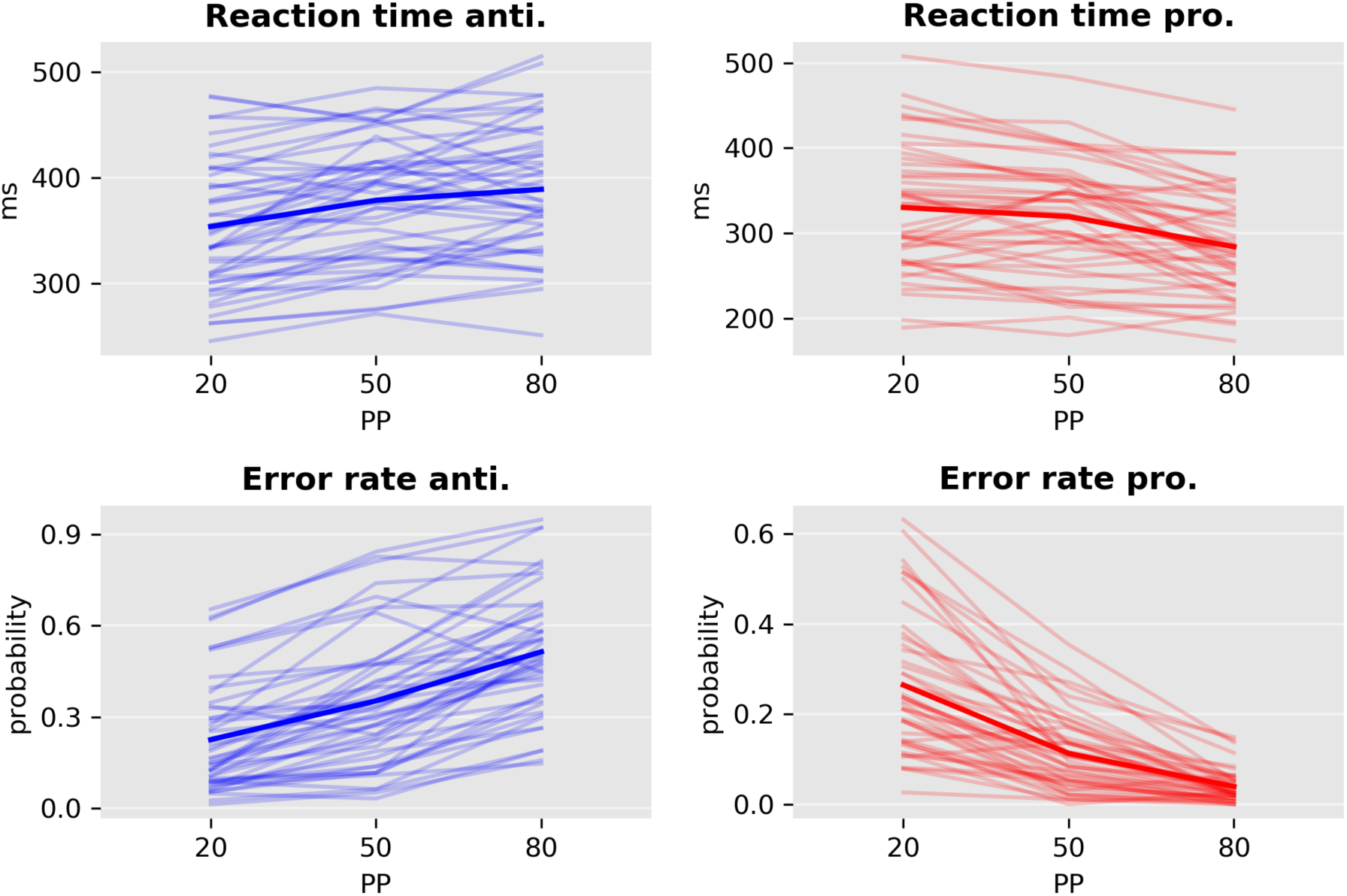
Reaction time and error rate in all conditions for each subject. Mean reaction times and error rates are displayed as solid lines.

